# Clinically Relevant Gene Editing in Hematopoietic Stem Cells for the Treatment of Pyruvate Kinase Deficiency Hemolytic Anemia

**DOI:** 10.1101/2021.01.14.426673

**Authors:** Sara Fañanas-Baquero, Oscar Quintana-Bustamante, Daniel P. Dever, Omaira Alberquilla, Rebeca Sanchez, Joab Camarena, Isabel Ojeda-Perez, Mercedes Dessy-Rodriguez, Rolf Turk, Mollie S. Schubert, Jose L. Lopez-Lorenzo, Paola Bianchi, Juan A. Bueren, Mark A. Behlke, Matthew Porteus, Jose-Carlos Segovia

**Affiliations:** Hematopoietic Innovative Therapies Division, Centro de Investigaciones Energéticas, Medioambientales y Tecnológicas (CIEMAT) and Centro de Investigación Biomédica en Red de Enfermedades Raras (CIBERER), Madrid, Spain; Unidad Mixta de Terapias Avanzadas. Instituto de Investigación Sanitaria Fundación Jiménez. (IIS-FJD, UAM). Madrid, Spain; Department of Pediatrics, Stanford University, Stanford, California, USA; Integrated DNA Technologies, Coralville, Iowa, USA; Hospital Universitario Fundación Jiménez Díaz. Instituto de Investigación Sanitaria Fundación Jiménez Díaz (IIS-FJD, UAM). Madrid, Spain; UOC Ematologia, Fisiopatologia delle Anemie, Fondazione IRCCS Ca’ Granda Ospedale Maggiore Policlinico Milano, Milan, Italy

## Abstract

Pyruvate Kinase Deficiency (PKD) is an autosomal recessive disorder caused by mutations in the *PKLR* gene, which constitutes the main cause of chronic non-spherocytic hemolytic anemia. PKD incidence is estimated in 1 in 20,000 people worldwide. The *PKLR* gene encodes for the erythroid pyruvate kinase protein (RPK) implicated in the last step of the anaerobic glycolysis in red blood cells. The defective enzyme fails to maintain normal erythrocyte ATP levels, producing severe hemolytic anemia, and can be fatal in severe patients. The only curative treatment for PKD is allogeneic hematopoietic stem and progenitor cells (HSPC) transplantation, so far. However, HSPC transplant is associated with a significant morbidity and mortality, especially in PKD patients. Here, we address the correction of PKD through precise gene editing at the *PKLR* endogenous locus to keep the tight regulation of RPK enzyme during erythropoiesis. We combined CRISPR/Cas9 system and rAAVs for donor matrix delivery to build an efficient and safe system to knock-in a therapeutic donor at the translation start site of the RPK isoform in human hematopoietic progenitors. Edited human hematopoietic progenitors efficiently reconstituted human hematopoiesis in primary and secondary immunodeficient recipient mice. Moreover, erythroid cells derived from edited PKD-HSPCs restored normal levels of ATP, demonstrating the restoration of RPK function in PKD erythropoiesis after gene editing. Our gene editing strategy may represent a lifelong therapy to restore RPK functionality in RBCs of patients and correct PKD.

**Single Sentence Summary:** Clinically relevant gene editing in hematopoietic stem cells for the treatment of Pyruvate Kinase Deficiency.

## INTRODUCTION

Pyruvate Kinase Deficiency is an inherited autosomal recessive metabolic disorder produced by mutations in the Liver and Erythroid Pyruvate Kinase Gene *(PKLR)*, which encodes for liver (LPK) and erythroid (RPK) pyruvate kinase proteins, expressed in liver and in red blood cells (RBCs) respectively. RPK is implicated in the last step of the anaerobic glycolysis pathway in RBCs. Glycolysis represents the main source of energy in RBCs. To date, more than 200 different mutations in the *PKLR* gene have been linked to PKD *(1, 2)*. PKD-causing mutations lead to a partial or total reduction in the RPK activity and the concomitant reduction in ATP levels, which favors RBC hemolysis and the consequent anemia. The disease becomes clinically relevant when the protein activity decreases below 25% of the normal activity in erythrocytes *(3)*. The most frequent clinical signs of the disease are mild to very severe anemia, reticulocytosis, splenomegaly and iron overload, implying that PKD might be life-threatening in severely affected patients *(4)*. PKD is considered the most common cause of Chronic Non-spherocytic Hemolytic Anemia (CNSHA). It shows a worldwide geographical distribution and the majority of the diagnosed patients are compound heterozygotes, as homozygous mutations are rare but very severe *(5)*. PKD is consider rare disease with an estimated prevalence of around 1:20,000*(1)(6)*, and higher in certain populations, such as the Amish community, as a result of a founder effect *(7, 8)*.

Treatments for PKD are mostly palliative and can help to improve the patient’s quality of life. The most extended is red blood cells transfusions, which can be occasional or very frequent, depending on the condition of the patient *(2)*. However, transfusions have related adverse effects, such as alloimmunization against donor blood cells and worsen of iron overload, which can result in important liver and heart organ damage *(2, 9)*. Iron chelation treatments has improved, but not solved, this life-threatening condition in PKD patients *(2)*. Spleen removal aims to prevent RBCs destruction, either reticulocytes or abnormal erythrocytes, and thus to increase the number of oxygen-transporting cells. Splenectomy does not arrest hemolysis but can increase hemoglobin (Hb) values up to 1-3 g/dL *(2)*. However, this treatment implies a risk of serious bacterial infections and an increased risk of venous thrombosis. Moreover, around 14% splenectomized PKD patients remain dependent on blood transfusion dependents after splenectomy. Currently, the only curative treatment for PKD is allogeneic hematopoietic stem cell transplantation (HSCT). However, it is not considered as routine treatment in PKD patients, due to the limitation of HLA compatible donors and the severe adverse effects, such as infections or development of graft-versus-host disease (GvHD), which can be particularly severe in PKD patients *(10–12)*.

Autologous HSCT of genetically corrected cells could overcome these limitations. This strategy has been used in several hematological genetic diseases *(13, 14)*, including hemoglobinopathies, *(15–17)(18)*, being already approved for clinical application (Zynteglo; http://shorturl.at/orMUY). We have recently developed a lentiviral vector to genetically correct PKD *(19)*, which has been granted orphan drug designation by the European and the American office regulators (EU/3/14/1330; FDA #DRU-2016-5168). This lentiviral-mediated gene therapy approach would offer a durable and curative clinical benefit with a single treatment, as shown by the preliminary results obtained in the first patient already infused with transduced autologous HSCs (NCT04105166) *(20)*.

Despite the promising results of conventional gene therapy, the ideal gene therapy approach should lead to the specific correction of the mutated gene, maintaining the endogenous regulation and eliminating the integration of exogenous DNA material elsewhere. Nuclease driven gene editing has emerged to allow it. This technology can be used for conducting precise double strand breaks (DSBs) and homologous recombination in the genome increasing up to 1000 times the previous efficacy of targeted modifications. A variety of genetic defects affecting non-hematopoietic tissues *(22–24)* and the hematopoietic system, such as X-linked SCID *(25)*, SCD *(26, 27)*, X-linked chronic granulomatous disease (X-CGD) *(28, 29)* or Fanconi anemia *(30, 31)* among others, have been successfully attempted by gene editing. Moreover, gene editing is already showing promising clinical results, as recently shown in hemoglobinopathies *(32–35)*. The improvement in the condition of two β-thalassemia and SCD patients enrolled in the CTX001® clinical trial *(36)*, pave the way to consider gene editing as a promising approach for RBC disorders, such as PKD. Previous studies performed in our laboratory demonstrated that gene editing in *PKLR* locus mediated by nucleases is feasible in iPSCs *(37)* and in HSPCs *(38)*, although the efficacy achieved was far from being clinically relevant. In this work, we have implemented the CRISPR/Cas9 nuclease system to induce DSBs at the transcription start site of *PKLR*, and recombinant adeno-associated vectors (rAAVs) to deliver a therapeutic DNA donor. We have been able to correct the phenotype of erythroid cells derived from PKD-HSPCs and to achieve clinically relevant levels of correction that envision the treatment of PKD patients by gene editing.

## RESULTS

### Design of the CRISPR/Cas9–AAV6-donor-transfer system to target *PKLR* gene in Hematopoietic Stem and Progenitor Cells

With the aim of developing a gene editing-based universal strategy for PKD patients, we followed a knock-in approach to insert a therapeutic donor at the translation start site of *PKLR* gene (Fig. S1 A and S2 A). To promote homologous directed repair (HDR) and favor integration of the therapeutic donor, we induced DSBs at *PKLR* transcription site by a specific CRISPR/Cas9 ribonucleoprotein (RNP). Therapeutic donor was delivered into target cells by adeno-associated vectorization as previously described*(26, 27, 39)*.

First, we designed different guide RNAs (gRNAs) to create DSBs around the start codon of the *PKLR* gene according to two main criteria, i) the highest on-target score possible (implying high specificity and less probable off-target [OT] activity) and, ii) the closest distance to *PKLR* starting site (Table S1). Efficacy of these gRNAs complexed with WT Sp. Cas9 protein (RNPs) was evaluated by TIDE assay in CB-CD34^+^ cells (Fig. 1A). SG1 produced the higher frequency of indels at the on-target site in human cells from healthy-donors (62.7±14.2%, Fig. 1A). To analyze the off-target (OT) activity, these guides were transfected into HEK293 cells stably expressing Cas9, to force OT effect, which allowed a stringent detection of OT by GUIDE-seq analysis *in vivo (40)*. OTs identified in this analysis are shown in Table S2. SG9 and SG10 presented reduced OT effect in comparison with the rest. Then, rhAmpSeq™ libraries were designed to analyze gene editing activity of the three most promising gRNAs, SG1 (the highest on-target activity in CB-CD34^+^ cells) and SG9 and SG10 (the lowest OT activity). Total gene editing modification at on-target sites of SG1, SG9 and SG10 was analysed by rhAmpSeq™ with or without dsODN donor (Fig. 1B and Fig S1B). Additionally, RNP complexes formed by either WT Cas9 or HiFi Cas9 activity were also evaluated in order to reduce OT effect *(27)*. SG1 gene editing activity without or with dsODN was higher than SG9 or SG10. Moreover, activity as RNP formed by HiFi Cas9 was not diminished (Fig 1B and Fig S1 B) that was confirmed by TIDE analyses (Fig. 1C). Lastly, OT effect of SG1 was analyzed through rhAmpSeq™ library of the top 49 OT sites in Jurkat cells (Fig. 1D). The percentage of gene modification in the most important OTs was reduced when HiFi Cas protein was used. In addition, a similar rhAmpSeq™ analysis was done in CB-CD34^+^ cells (Fig. 1E). Gene editing of top ten OT sites in hCD34^+^ cells were below 0.1% total gene editing when HiFi Cas9 was used. Altogether, these analyses demonstrated that HiFi-Cas9/SG1-RNP promotes significant levels of perfect HDR in the on-target site, with a minimal effect in the potential off-targets, confirming the safety of the use of this RNP in the *PKLR* locus.

**Figure 1.**
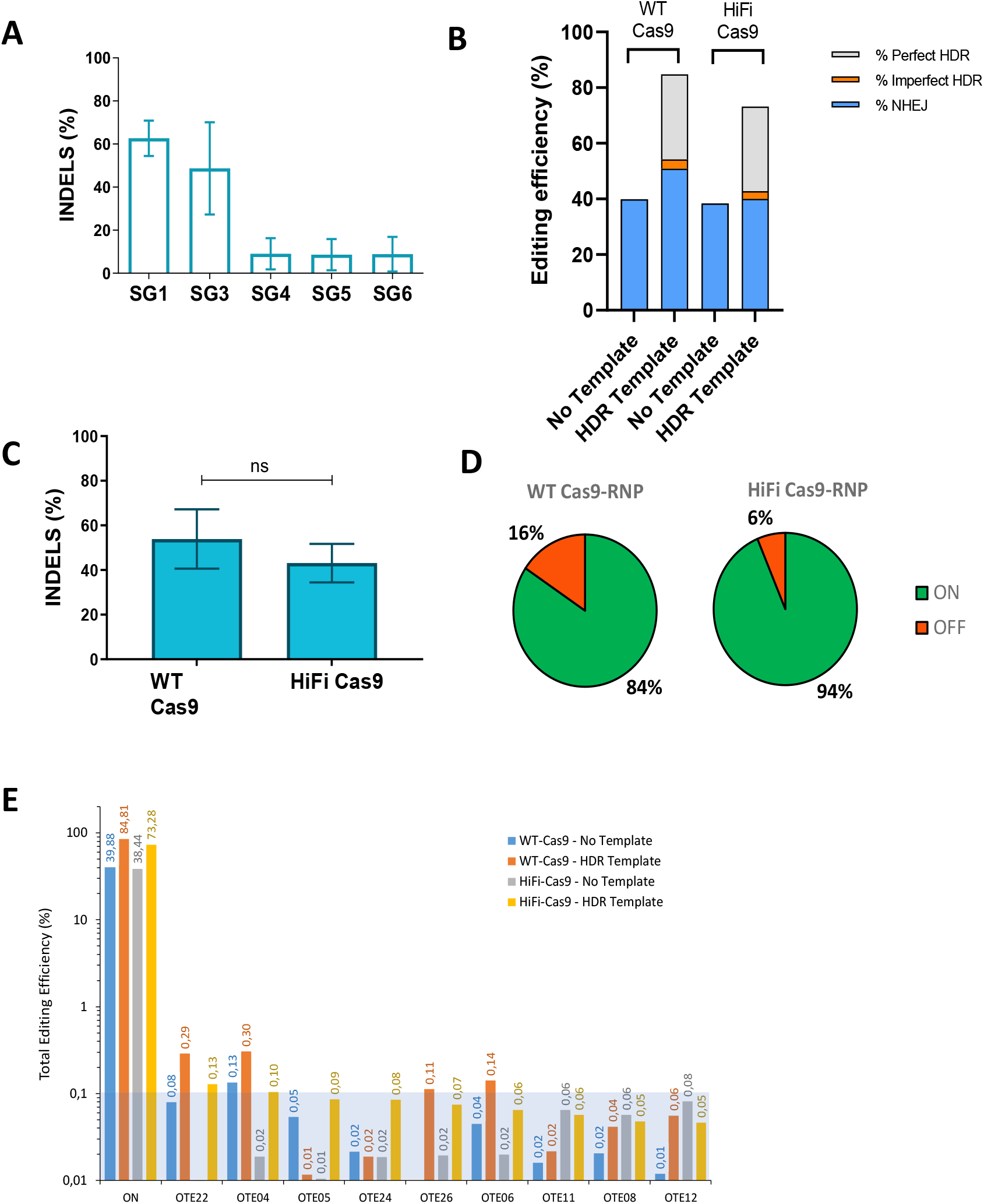
Selected sgRNA displays high on target efficiency and very low off-target incidence. **A)** Generation of insertions and Deletions (InDels) in CB-CD34^+^ cells with different guides selected around the transcription starting site of the *RPK* cDNA; Data are represented as mean ± SD; N=3; **B)** Analyses of the HDR vs NHEJ in samples edited with SG1, complexed with either WT Cas9 or HiFi Cas9. Blue, % of NHEJ repair, Gray, % of perfect HDR, Orange, % of imperfect HDR. **C)** Comparison of the generation of InDels in CD34^+^ cells by SG1 electroporated as RNP using, wild-type or HiFi Cas9 protein; Data are represented as mean ± SD; N=3; **D)** On-target and Off-target frequency estimated by GUIDE-seq using either wild-type or HiFi Cas9 in the RNP complex; **E)** Analyses of total gene editing efficiency by Cas9 HiFi+SG1±HDR-SG1 template in the top ten sites ranked high-low. In orange and yellow bars, gene editing caused by SG1 complexed with WT Cas9 or HiFi Cas9, respectively, in the absence of the specific HDR template. In blue and grey bars, the gene editing caused by SG1 complexed with WT Cas9 or HiFi Cas9, respectively, in the presence of the specific HDR template. Blue square indicates the limit of detection of the assay (<0.1%). Significance was analyzed by non-parametric two tailed Mann Whitney’s test.

The design of the donor sequence to be packaged in AAV viral vector was conducted considering the selected SG1. Two donors were designed (Fig.S2 A), a reporter donor, carrying a turbo-GFP cDNA under the regulation of Ubiquitin C promoter (UBC) and a therapeutic donor, which carried the corrective sequence (coRPK cDNA), without any exogenous promoter, to allow the endogenous *PKLR* promoter to drive the expression of the therapeutic coRPK cDNA (see details in Materials and Methods). Donor sequences were flanked left and right homology arms, with AAV ITRs and packaged into AAV-6 serotype.

### Homology Directed Repair in the *PKLR* locus is effective in an erythroleukemic cell line and in human hematopoietic progenitors

Gene editing tools were validated in K562 human erythroleukemia cells. Cells were electroporated with the SG1-RNP and transduced with rAAVs carrying either the reporter or the therapeutic donor, at a viral concentration of 1×10^4^ genome copies per microliter (gc/µl). Cells transduced with the reporter donor were visualized by fluorescence microscopy 5 days post-transduction (dpt). Gene targeting efficiency of around 30% was estimated by immunofluorescence (Fig.S2 B). DNA from samples edited with the therapeutic vector was analyzed by specific PCRs amplifying the genomic junctions between the endogenous and exogenous DNA both at 5’ and 3’ ends to verify the correct integration of the transgene. Results confirmed the HDR of the therapeutic coRPK donor at the on-target site (Fig.S2 C-D). Furthermore, we quantified the expression of the endogenous RPK and therapeutic coRPK transcripts through qRT-PCR. Transduced and untransduced cells expressed the WT endogenous RPK mRNA (Fig. S2 E-F). However, coRPK transcripts were exclusively detected in K562 cells transduced with the therapeutic vector.

Next, we assessed the targeting efficiency in human HSPCs. CB-CD34^+^ cells were nucleofected with *PKLR* SG1 RNP and then transduced with either of the two AAV donors. Forty-eight hours after nucleofection, clonogenic potential of edited cells was assessed in semisolid cultures by colony-forming units assay. No differences in the number or in the hematopoietic lineage distribution were found when cells were edited with either donor (Fig.2A). GFP^+^ colonies visualized in colonies from CD34^+^ cells transduced with the reporter donor (Fig.S3 A), reached values up to 38 % GFP^+^ CFUs (21.1±17.44%) (Fig.S3 B). Correct integration of the therapeutic donor was verified by PCR in individually picked colonies. In a representative agarose gel (Fig.2 B), the specific 852bp (right arm) and 516bp (left arm) bands appeared in CFUs derived from edited CB-hCD34^+^ cells. In addition, specific integration of the reporter donor was also confirmed by PCR in CFUs (Fig.S3 C). PCR amplicons of individual CFUs were Sanger sequenced and the results confirmed the correct integration of the reporter and therapeutic donors in human progenitor cells (Fig.2 C and Fig. S 3D). In five different experiments, the percentage of positive colonies that displayed correct integration of the reporter and the therapeutic donor sequences in 5’ end (LHA) and 3’ end (RHA) was 35.5±4.7% and 38.5±7.4%, respectively (Fig.2 D). Altogether, results evidence the efficient knock-in of the desired donor at the starting site of *PKLR* gene in human hematopoietic progenitor cells.

**Figure 2.**
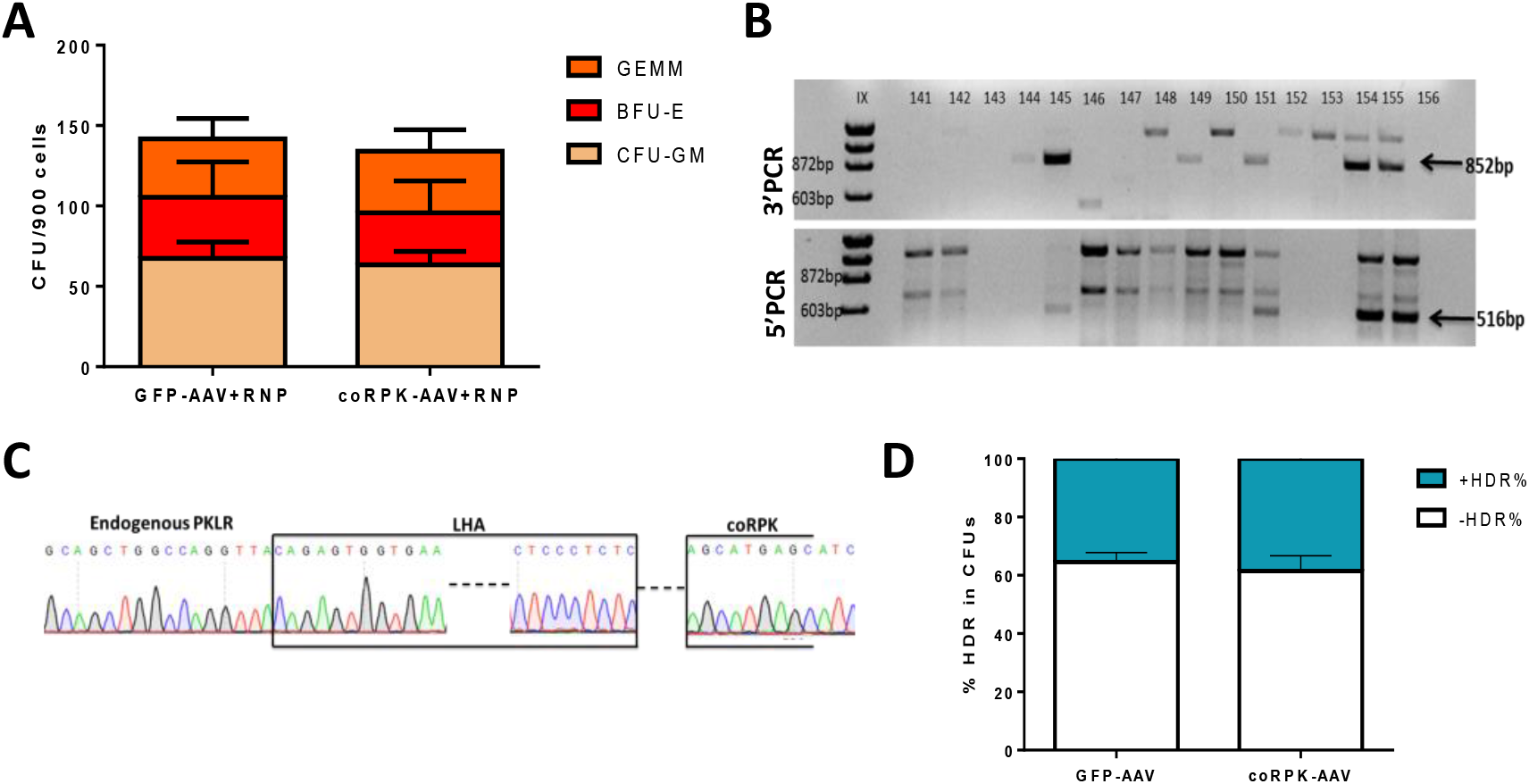
RNP plus AAV6 efficient gene targeting in human hematopoietic progenitor cells. **A)** Colony Forming Unit number and distribution is maintained when cells are transduced with either of the donor AAVs in 5 independent experiments. GEMM, Granulocyte/erythrocyte/monocyte/megakaryocyte progenitors; BFU-E, Burst Forming Unit-Erythroid; CFU-GM, Colony Forming Unit-Erythroid; **B)** Representative agarose gel of the PCR analyses of the specific integration of coRPK in individually picked colonies by amplification of the right and the left homology arms (RHA, 852bp and LHA, 516bp, respectively) of the donor construct; **C)** Representative sanger sequencing of one of the positive amplicons; **D)** Percentage of correct HDR in CFUs assessed by PCR when cells were edited with GFP-AAV or coRPK-AAV and RNP complex. Mean±SD is indicated.

### *PKLR* locus is efficiently targeted in long-term hematopoietic repopulating stem cells

To test if the more primitive hematopoietic stem cells (HSCs) were also targeted using the described strategy, new CB-hCD34^+^ cells from healthy donors were nucleofected with RNP-SG1 and transduced with either reporter or therapeutic donor. Transduction efficiency assessed by flow cytometry 48hpt was 20% GFP^+^ in CD34^+^ cells. Within them, 5-10% of CD34^+^CD38^-^CD90^+^ primitive HSCs were GFP^+^ (Fig. 3A). To evaluate the editing efficiency in hematopoietic repopulating cells, 8×10^5^-1×10^6^ edited cells were transplanted into sublethally irradiated immunodeficient NSG mice. The follow-up of human hematopoietic engraftment at 1 and 3 months post-transplant demonstrated that gene edited cells were able to completely repopulate immunodeficient recipients (Fig. 3B). Ninety days post-transplant, mice transplanted with GFP-AAV edited cells reached 87.27±8.42% human chimerism, and mice transplanted with coRPK-AAV edited cells presented similar levels (87.13±5.86%) with no signs of toxicity related to the gene editing process when compared with laboratory historic data of transplants with non-manipulated cells (Fig. 3B). Percentage of GFP^+^ cells within the human compartment in the BM of the recipients transplanted with cells transduced with the AAV reporter donor was 2.29±1.89% at 1mpt and 0.34±0.41% at 3mpt (Fig. 3C and Fig. S4A). Percentage of GFP^+^ cells within the hCD34^+^ compartment analyzed 3mpt was 2.2±3.63% (Fig.3 D). Multilineage differentiation in bone marrow cells was also investigated using antibodies against hCD34 for HSPCs, hCD33 for myeloid cells and hCD19 for lymphoid cells (Fig. 3E-F). Edited cells were found within the three human hematopoietic subpopulations (Fig. S4B), confirming the gene editing of the HSCs capable of generating myeloid, lymphoid and progenitor cells. To confirm the long-term repopulating capacity of edited cells, BM cells from primary recipient were transplanted into secondary recipients. FACS analyses of hCD45^+^ cells in BM of secondary recipients showed that edited cells were able to efficiently repopulate secondary recipients. In the case of recipients infused with the GFP-edited cells 16.87±15.74% of mouse cells were hCD45^+^ cells, and a similar value of 17.77±12.60% hCD45^+^ cells was observed in the BM of secondary mice infused with cells edited with therapeutic donor. In all instance, a multilineage differentiation engraftment was observed (Fig. 4A-C). Human CD45^+^GFP^+^ and CD34^+^GFP^+^ cells were detected in secondary recipients as well (Fig. 4D-E), and the percentage of GFP^+^ cells within the human population was maintained along the time (3.27±5.66% total human cells and 0.82±1.41% human progenitor cells 3mpt). Human erythroid compartment, identified as mTer119^-^ hCD235a^+^hCD71^+/-^ was also analyzed. GFP^+^ cells were found within the human erythroid population, and erythroid differentiation was not affected when the therapeutic donor was used (Fig. 4F-G). BM cells from secondary recipients, which were transplanted with HSCs edited with the SG1-RNP and the therapeutic donor, were analyzed by specific PCR, confirming the specific knock-in of the coRPK cDNA (Fig.4 H). Taken together, these results demonstrate the efficient knock-in gene editing protocol in HSCs capable of repopulating in the long term the hematopoiesis of immunodeficient NSG mice.

**Figure 3.**
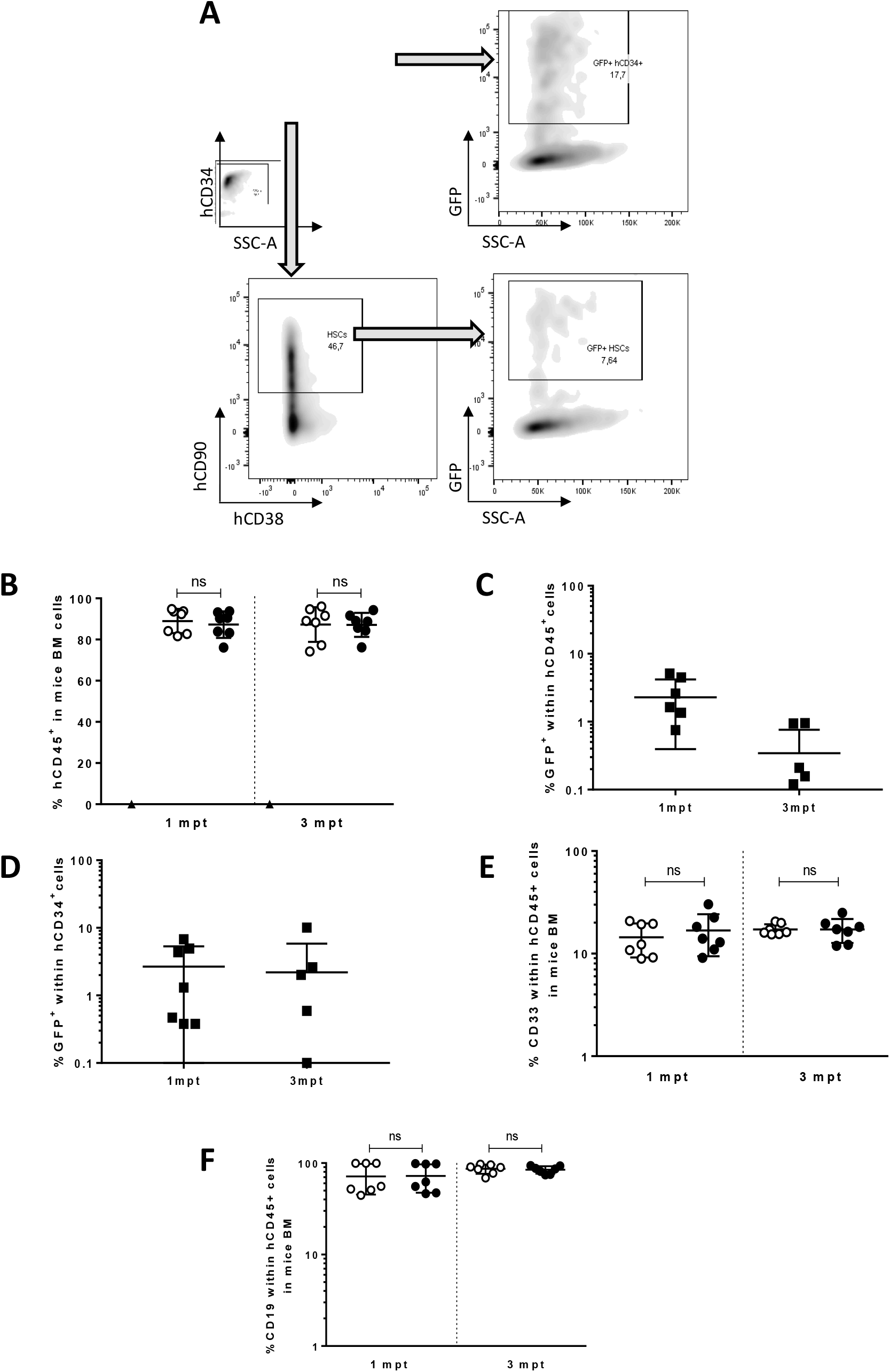
Targeted human hematopoietic stem cells engrafted in immunodeficient mice. **A)** FACS analysis 48hpt of hCD34^+^ cells electroporated with the specific RNP and transduced with GFP-AAV donor. GFP% within the total hCD34^+^ cells and within the most primitive stem cells, marked as hCD34^+^hCD38^-^hCD90^+^; **B)** Percentage of human cells in the BM of immunodeficient NSG mice transplanted with GFP-AAV-(white dots) or therapeutic-AAV-(black dots) edited cells, one and three months post-transplant. Triangle dot corresponds to one control animal transplanted with irradiated-hCD34^-^ cells; Data are represented as mean ± SD (n = 7 in each group); **C)** Percentage of GFP^+^ cells within the human population (hCD45^+^) one and three months post-transplant; Data are represented as mean ± SD (n = 7); **D)** Percentage of GFP^+^ cells within the human progenitor (hCD34^+^) cells one and three months post-transplant; Data are represented as mean ± SD (n = 7); **E)** Percentage of the hCD33^+^ cells within the human population (hCD45^+^) in mice transplanted with cells edited with reporter (white dots) or therapeutic donor (black dots). Data are represented as mean ± SD (n = 7 in each group); **F)** Percentage of the hCD19^+^ cells within the human population in mice transplanted with cells edited with reporter (white dots) or therapeutic donor (black dots). Data are represented as mean ± SD (n = 7 in each group); A two-way ANOVA was performed followed by Tukey’s post hoc test. ns= not significant.

**Figure 4.**
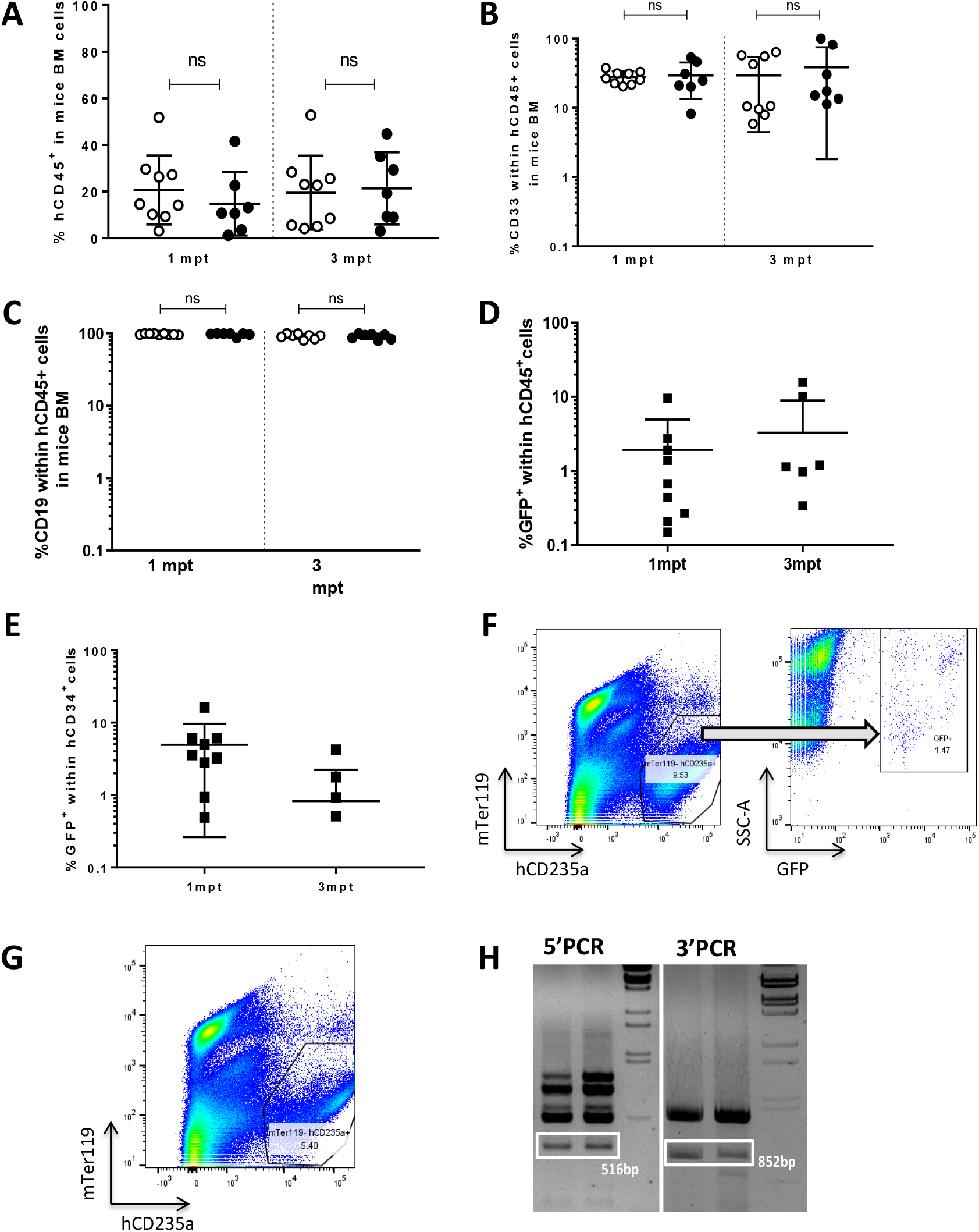
Gene edited HSC are able to efficiently engraft and differentiate in secondary transplanted immunodeficient mice. **A)** Percentage of human CD45^+^ cells in immunodeficient NSG animals transplanted with total BM of primary transplanted animals with either GFP-AAV-(white dots; n=9) or therapeutic-AAV-(black dots; n=7) edited cells analyzed one and 3 months post-transplant; Data are represented as mean ± SD; **B)** Percentage of the hCD33^+^ cells within the human population (hCD45^+^) in mice transplanted with cells edited with reporter (white dots; n=9) or therapeutic donor (black dots; n=7). Data are represented as mean ± SD; **C)** Percentage of the hCD19^+^ cells within the human population in mice transplanted with cells edited with reporter (white dots; n=9) or therapeutic donor (black dots; n=7). Data are represented as mean ± SD; **D)** Percentage of GFP^+^ cells within the human population (hCD45^+^) one and three months post-transplant; Data are represented as mean ± SD; **E)** Percentage of GFP^+^ cells within the human progenitor (CD34^+^) cells one and three months post-transplant; **F)** Representative dot-plot of human erythroid differentiation in secondary transplanted recipients of GFP-AAV-edited cells. Left, CD235a vs mTer119 expression to identify human erythroid cells. Right, GFP expression within human erythroid population (CD235a^+^); **G)** Representative dot-plot of human erythroid differentiation in secondary transplanted recipients of therapeutic-AAV-edited cells. Left, CD235a vs CD71 expression in human cells. Basophilic erythroblasts (gate A) and polychromatic plus orthochromatic erythroblasts (gate B) are shown; **H)** Representative agarose gel of PCR analysis of the 5’ (LHA, 516 bp) and the 3’ (RHA, 852 bp) sequences in two different mice’s total BM cells, demonstrating correct integration of the donor fragment. A two-way ANOVA was performed followed by Tukey’s post hoc test. ns= not significant.

### Optimized gene targeting conditions in hCD34^+^ cells allow clinically relevant efficacies

Unpublished data with lentiviral vectors from our laboratory point-out that 25-30% corrected cells are necessary to observe a clinical improvement in PKD mouse model (S. Navarro et al, submitted). In order to assess the therapeutic potential of our gene editing system, we intended to maximize the efficacy of the transduction to reach the therapeutic limit. Doses of the reporter AAV ranging from 1×10^1^ to 1×10^5^gc/µl were tested (Fig. 5A). Vector concentrations of 2.5×10^4^ and 5×10^4^gc/µl displayed enhanced editing efficacies of 25.35±0.35% and 33.52±15.09% GFP^+^ cells, respectively. Remarkably, 14.49±8.93% of the hCD34^+^hCD38^-^hCD90^+^ cells were GFP^+^ when samples were transduced with 5×10^4^gc/µl AAV (Fig. 5A and Fig. S5A). No significant differences in cell viability of the cell culture were observed when higher and more efficient viral vector concentrations were used (5×10^4^gc/µl) in comparison with previously used (1×10^4^gc/µl) (Fig. 5B). Gene targeting efficiency assessed in the BM of mice transplanted with edited cells was definitely increased when the higher AAV dose was used (20.38±11.76% in contrast with 0.34±0.41% achieved with 10^4^gc/µl) (Fig. 5C). These data reveal that levels of gene editing obtained were in the range of those required to have clinical benefit to correct PKD.

**Figure 5.**
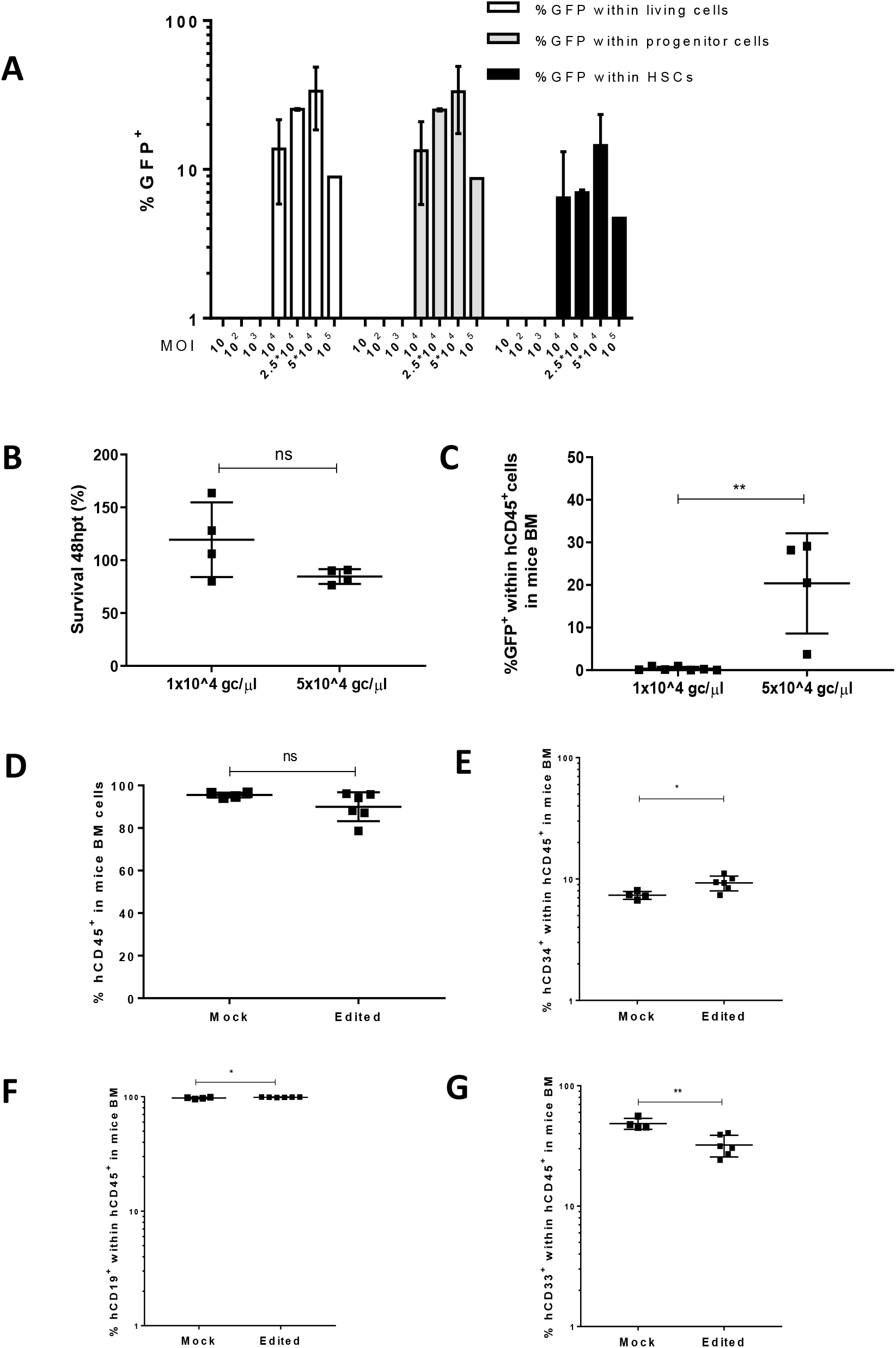
Optimization of the gene editing protocol is able to restore the energetic balance in PKD-HSPCs. **A)** Percentage of GFP cells in different subpopulations (living cells, progenitor cells [hCD34+] and HSCs [hCD34^+^hCD38^-^hCD90^+^] 48hpt using different MOIs ranging from 10 to 100000gc/µl of the reporter donor. **B)** Viability in HSPCs is not affected when increasing the vector dose 5 times. **C) %** GFP is significantly higher in mice receiving HSPCs edited with 5×10^4^gc/µl GFP-AAV when compared with mice receiving cells edited with 10^4^gc/µl. **D)** Percentage of human CD45^+^ cells in immunodeficient NBSGW animals transplanted with mPB CD34^+^ cells from a PKD patient, previously edited with therapeutic-AAV-edited cells, analyzed 2 months post-transplant; **E)** Percentage of the hCD34^+^cells within the human population(hCD45^+^) in animals engrafted with PKD edited human cells; **F)** Percentage of the hCD19^+^cells within the human population (hCD45^+^)^+^) in animals engrafted with PKD edited human cells; **G)** Percentage of the hCD33^+^cells within the human population (hCD45^+^) in animals engrafted with PKD edited human cells. Kruskal-Wallis multiple comparison test was performed; P value is indicated in the figure. ns=not significant

### ATP deficiency is corrected after *PKLR* gene editing in PKD-HSPCs

Optimized editing conditions were then tested in BM-CD34^+^ cells from three PKD patients and from mobilized PB-CD34^+^ cells from 1 PKD patient, carrying different mutations in the *PKLR* gene (Table S4). Cells were pre-stimulated for 48 hours, nucleofected and then transduced with the rAAVs, or submitted to the same protocol without the gRNAs and the therapeutic rAAV. Healthy donor-CD34^+^ cells were sham nucleofected and transduced as controls. Twenty-four hours after the gene editing procedure, cells were collected and transferred to an *in vitro* erythroid differentiation protocol. Edited cells from one of the patient samples were transplanted into immunodeficient NBSGW mice to facilitate the analysis of human erythroid subpopulations. Sixty days after transplantation, levels of human hematopoietic chimerism were analyzed in mice BM. Both groups of mice (transplanted with mock or edited cells) showed high percentages of human engraftment (95.5±1.11% and 89.98±6.78% hCD45^+^ cells, respectively) (Fig. 5D) and no differences in the multilineage engraftment were observed between mock and edited cells (Fig. 5E-G). Specific integration of coRPK was assessed in pre-transplant samples, in *in vitro* erythroid differentiated samples and in the BM of the mice transplanted with mock or edited cells. In all cases, only edited cells showed the specific editing in 3’ and 5’ junctions (Fig. 6A). Expected bands were also found in hCD19^+^ and hCD33^+^ sorted populations from the transplanted NBSGW mice BM (Fig. S5 C), confirming that editing had occurred in multipotent HSCs. In parallel, erythroid differentiation process was evaluated along time by FACS. As shown in Fig S5 B, no significant differences between healthy and PKD donor samples were observed. In order to assess the percentage of gene targeting in the engrafted cells, we performed a CFU assay with sorted BM hCD34^+^ cells. Up to 61 individual colonies from 5 different transplanted mice were analyzed. Among colonies positive for the GAPDH housekeeping PCR (51 colonies in total), 12 presented the expected integration of coRPK sequences both at 5’ and 3’ junctions, indicating an efficiency of 25.24±7.28% HDR in PKD patient’s HSCs (Fig. 6B). As expected, no colonies from mock edited colonies showed coRPK integration (Fig. 6B).

**Figure 6.**
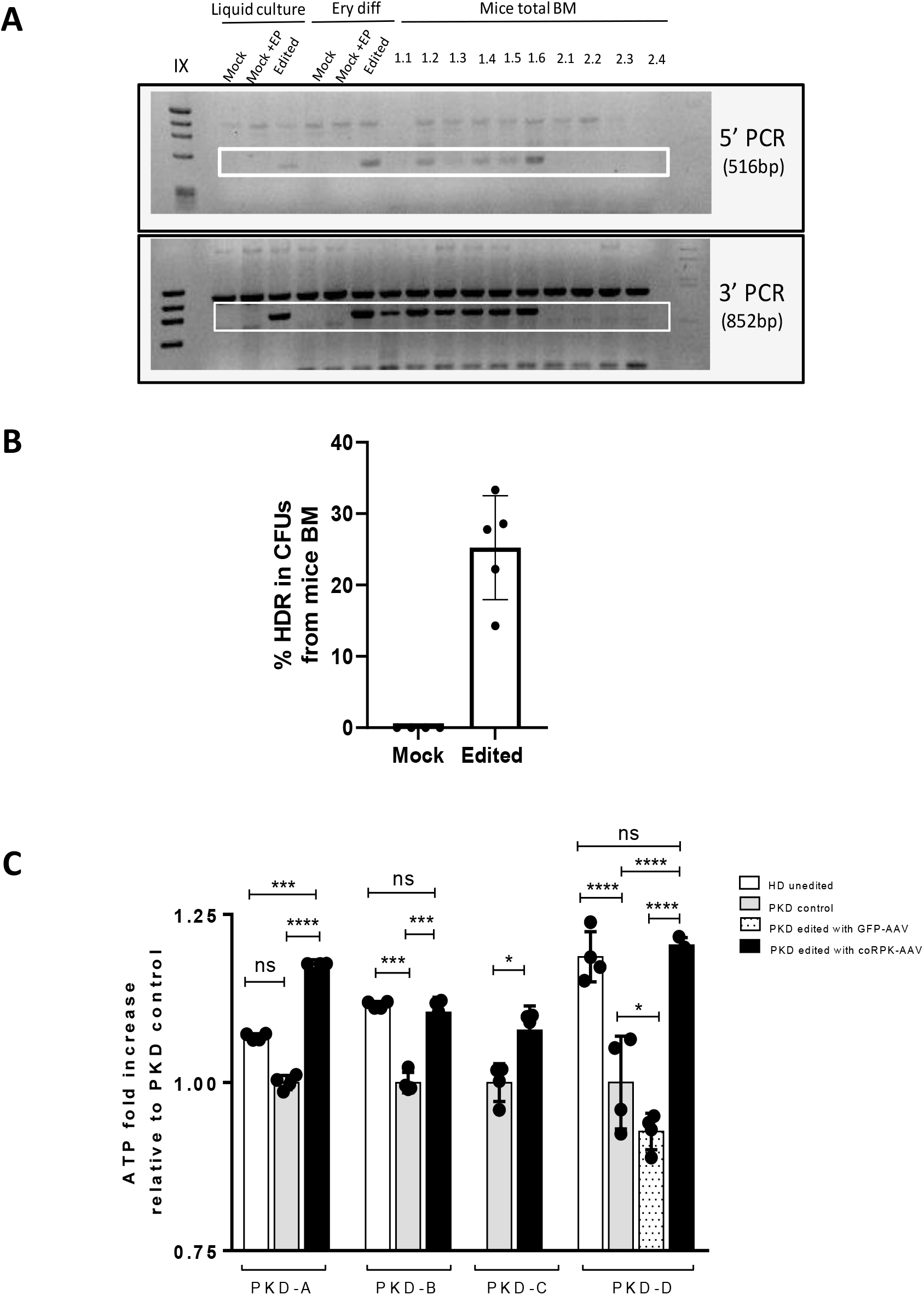
Editing in engraftable PKD-HSPCs is feasible. **A)** Representative agarose gel of 5’ and 3’ PCRs in samples pre-transplant (liquid culture), after erythroid differentiation (Ery diff), in mice receiving edited cells (1.1, 1.2, 1.3, 1.4, 1.5 and 1.6) and in mice receiving unedited cells (2.1, 2.2, 2.3 and 2.4). Mock: unedited cells; Mock+EP: unedited electroporated cells; coRPK: cells edited with therapeutic donor; Squares identify amplification of the band showing homologous directed integration; **B)** hCD34+ cells were sorted from the total BM of NBSGW mice transplanted with PKD-D edited HSPCs. Sorted cells were cultured in methylcellulose and the percentage of HDR was assessed by specific PCR in 5’ and 3’ junctions. Data are represented as mean ± SD; **C)** ATP quantification in *in vitro* differentiated erythroid cells obtained from CD34^+^ cells of four PKD patients, previously edited with the GFP-AAV or the therapeutic-AAV donors. Three different healthy donors were also in vitro differentiated and analyzed. Data are relativized to unedited erythroid PKD cells. HD unedited: HSPCs from a healthy donor that had undergone *in vitro* erythroid differentiation; PKD unedited: HSPCs from a PKD patient that had undergone *in vitro* erythroid differentiation; PKD GFP/coRPK: HSPCs from a PKD patient that had undergone gene editing with GFP-AAV or coRPK-AAV and subsequent *in vitro* erythroid differentiation. Individual dots are replicates of the luminescence measurement and bars represent the media of the replicates. Kruskal-Wallis multiple comparison test was performed; P value is indicated in the figure. The significance was represented by P-values: *P<0.05, **P<0.01, ***P<0.005 and ****P<0.001. ns=not significant.

Finally, a functional analysis based in the quantification of ATP in *in vitro* differentiated erythroid cells was performed in the four edited samples. Unedited PKD cells or cells edited with the reporter vector produced low levels of ATP. However, erythroid cells that arise from PKD-CD34^+^ cells that had undergone gene editing with the RNP and therapeutic donor were able to restore the ATP levels similarly to those determined by HD cells (Fig. 6C). Altogether, these results demonstrate the clinically relevant application of CRISPR/Cas9 and AAV-based gene editing to address the treatment of PKD patients.

## DISCUSSION

PKD is nowadays considered a good candidate for gene therapy, supported by key factors, such as the monogenic origin of the disease and the confirmation of the efficacy of allogenic HSCT as a curative treatment for some PKD patients *(12)*. Additionally, as initially reported by *García-Gómez et al (19)*, the correction of the *PKLR* gene defect by means of lentiviral-mediated gene therapy is feasible and restores RPK functionality. In fact, an international multicentric clinical trial for the treatment of PKD patients is currently on-going (NCT04105166), and has already shown first evidences of therapeutic efficacy *(20)*. Despite this promising therapy for PKD patients, the emergence of programmable nucleases has revolutionized the gene therapy field, making specific driven integration gene therapy an applicable clinical option *(36)*. We have previously shown that gene editing in *PKLR* gene is achievable in iPSCs *(37)* and in HSPCs *(38)*, showing correction of the PKD phenotype, although the achieved efficiencies were far from a potential clinical application*(3, 19)*.

The feasibility of knocking-in a cDNA immediately after the start codon of the gene has been recently demonstrated *(26, 42, 43)*. This approach allows the restoration of the specific gene functionality without compromising the endogenous regulatory control of the gene expression in all possible mutations affecting the open reading frame. In our case, this strategy would be applicable to most PKD patients, with the exception of the ones carrying mutations in the promoter or regulatory regions (only 1.1% of mutations happen to be in promoter sequences *(9)*). The developed system does not require selection. The genetic tools are easily designed and synthesized, and showed high levels of efficiency. The best RNP cutting in the start codon of *PKLR* gene was selected following two main criteria: the highest ON and lowest OT profiles. SG1-RNP produced DSBs above 62.7±14.2% in CB-hCD34^+^ cells while maintaining its cleavage in the top 10 OTs under 0.1% assessed by GUIDE Seq, when *PKLR* SG1 guide RNA was complexed with HiFi Cas9, an engineered version of Sp.Cas9. To assess the safety of the designed *PKLR* guides, we have used a robust technique, such as GUIDE-seq, which ensures the safety of the *PKLR* SG1 regarding its OT effect. This strict requirement of little OT activity is crucial when a multiclonal cell population is targeted for clinical uses, as it is the case for HSPCs gene therapy. Additionally, to deliver HDR donors, we selected the AAV6 platform, which is been widely used to edit human HSPCs. A therapeutic AAV6 donor carrying the coRPK cDNA and no promoter regions was synthesized (therapeutic donor). In parallel, a reporter donor was also generated, and both therapeutic and reporter donors’ cDNA were diverged to prevent re-cutting of the SG1-RNP after gene editing has occurred. The combination of SG1-RNP and either therapeutic or reporter donor AAV6 boosted the efficacy of HDR at *PKLR* locus in human hematopoietic cells. No toxic effects linked to the protocol were found in CFUs, and indeed, when transduced with reporter donor, 21.1±17.44% CFUs were GFP^+^ (Fig. S3B). Additionally, 35.5±4.66% and 38.5±7.38% of the CFUs presented the specific integration of the reporter donor or the therapeutic donor, respectively (Fig. 2D). The discrepancy between the fluorescence analyses and the molecular characterization could be attributed to differences in the sensitivity in the GFP^+^ quantification, since the monoallelic integration could minimize the fluorescence intensity. Furthermore, edited cells, when transplanted into immunodeficient mice, were able to repopulate very efficiently primary and secondary recipients (Fig.3-4). Although edited cells were able to give rise to different hematopoietic lineages, the percentage of gene editing within the human cells in mice was below 5% (Fig. 3C and Fig. 4D). Consequently, we attempted some optimizations of the protocol and we observed that increasing 5-fold the vector concentration allowed us to achieve higher percentages of gene editing with the reporter donor in CFUs and in xenotransplanted mice, without compromising human cell viability or stem potential (Fig. 5A-C). More importantly, when we applied these optimizations to PKD patients’ HSPCs we observed that edited PKD-HSPCs were able to efficiently reconstitute the BM of NBSGW mice (Fig. 5D-G) and coRPK integration was observed in the mice transplanted with edited cells (Fig. 6A). We corroborated 25.24±7.28% HDR by specific in-out PCRs in hCD34^+^ cells from the BM of transplanted mice (Fig. 6B), a value that is within the therapeutic window for PKD correction. Furthermore, after *in vitro* differentiating edited PKD-HSPCs towards the erythroid lineage, we observed a restoration in the functionality of RPK protein to almost healthy levels, measured by ATP production in four different patients (Fig. 6C).

This knock-in strategy represents the most promising gene editing approach in *PKLR* gene so far. It works both *in vitro* and *in vivo*. Optimizations in pre-stimulation time and vector concentration have improved the outcome of the protocol, but transduction efficiency in the most primitive compartment (HSCs) is still in the limits of the therapeutic application. We have considered the use of different editing enhancers, such us small molecules or specific microRNAs, to enhance the knock-in in HSPCs. In previous gene editing experiments focused on the knock-in in the second intron of the gene *(38)*, dimethyl prostaglandin E2 did not enhance the procedure, but it may be worth testing it in this non-selection system since it was reported to enhance gene editing in engraftable HSCs in similar contexts *(25)*. Besides, other molecules such as Scr7 have been shown to affect NHEJ pathway by the inhibition of DNA ligase IV, a key enzyme for this pathway. Thus, the downregulation of NHEJ increased the efficiency of HDR in mammalian cells *(44, 45)*. Other groups have reported the use of SR-1 molecule as an HDR pathway enhancer in TALEN and CRISPR/Cas9-mediated editing systems *(46)*. Aside from small molecules, it has been recently reported that inhibition of p53 increases the rate of HDR *(47)*. Furthermore, *Schiroli et al* reported that AAV6-mediated gene editing aggravates p53 activation and delays HSPCs proliferation *(39)*, which can explain our limited levels of HDR in long-term HSCs. They also claimed that transient p53 inhibition (during the first 24 hours post-editing) alleviated repopulating defects in edited HSPCs and did not lead to any chromosomal aberrations. Either way, inhibiting p53 is a risky approach, since it is widely known that stable inactivation of p53 pathway can lead to development of malignancies. However, a recent study in mice reported no increases in mutational load upon stable p53 genetic inactivation in HSCs *(48)*, opening the door to the possible regulation of p53 during the gene editing procedure in order to increase the yield of edited HSPCs that could engraft in patients and ensure rapid establishment of therapeutic benefit.

In summary, the present study demonstrates that a gene editing approach based on RNP electroporation and donor rAAV transduction in PKD HSCs is safe and efficient to correct PKD. Moreover, the levels of HSPCs-gene editing achieved with the proposed strategy are in the range required to be of clinically relevance for the treatment of PKD patients.

## MATERIALS AND METHODS

### Human cells

K562 cell line (chronic myelogenous leukemia; ATCC: CCL-243) was cultured in Iscove’s modified Dulbecco’s medium (IMDM; Gibco), 20% HyClone™ Fetal Bovine Serum (FBS, GE Healthcare) and 1% penicillin/streptomycin (P/S) solution (Gibco). Cells were maintained at 5×10^5^-1×10^6^ cell/mL.

Umbilical cord blood samples (CB) from healthy donors were provided by *Centro de Transfusiones de la Comunidad de Madrid* and samples from PKD patients were provided by *Hospital Universitario Fundación Jiménez Díaz, Hospital Infantil Universitario Niño Jesús* and *Ospedale Maggiore Policlinico*. All samples were collected under written consent from the donors and *Centro de Transfusiones de la Comunidad de Madrid’s* institutional review board agreement (number PKDefin [SAF2017-84248-P]). Mononuclear cells were obtained by Ficoll-Paque PLUS (GE Healthcare) density gradient isolation according to manufacturer’s recommendations. Purified CD34^+^ cells were obtained by immunoselection using the CD34 Micro-Bead kit (MACS, Miltenyi Biotech). Magnetically-labelled cells were selected with LS and MS columns sequentially in QuadroMACSTM Separator (Milteny Biotech) following manufacturer’s instructions. Purified cells were kept frozen or used fresh in further experiments.

Cells were grown in StemSpan (StemCell Technologies) supplemented with 0.5% P/S, 100 ng/ml human Stem Cell Factor (SCF), 100 ng/ml human thrombopoietin (TPO), 100 ng/ml human FMS-like tyrosine kinase 3 ligand (Flt3), 100ng/ml human interleukin 6 (hIL-6) (all obtained from EuroBiosciences) and 35nM UM171 molecule (Stem Cell Technologies). Cells were cultured under normoxic conditions: 37°C, 21% O_2_, 5% CO_2_ and 95% relative humidity.

### Guide RNAs

The design of the different guide RNAs to introduce DSBs in the genomic sites of interest was performed using the different website tools available for that purpose, such as Dr. Zhang’s lab tool (https://zlab.bio/guide-design-resources) or Integrated DNA Technologies (IDT) website (https://eu.idtdna.com/site/order/designtool/index/CRISPR_SEQUENCE).

Additionally, the activity of the designed guide was assessed through calculating the insertion-deletion (Indel) frequencies using the TIDE software (https://tide.deskgen.com/). PCR of genomic DNA extracted 3 days after nucleofection using NucleoSpin Tissue kit (Macherey-Nagel) was performed using specific primers and Sanger sequenced (Stabvida, Caparica, Portugal). Unedited cells were always used as a negative control for calculating INDEL frequencies with TIDE. Sanger sequencing was done with ATG TIDE F 5’-CCTGCTCCCTGGATTCACTA-3’ and ATG TIDE R 5’-TTTAACACACGGGAGGCTCT-3’ primers.

### rAAV/Cas9 gene editing

To assemble ribonucleoproteins (RNP), 6µg Alt-R® S.p. Cas9 Nuclease V3 (IDT) was combined with 3.2µg synthetic sgRNAs (Synthego) at RT for 10 min immediately prior to be used.

Human K562 cells were nucleofected with already complexed RNP using the SF Cell Line 4D Nucleofector X Kit for Amaxa 4D device (Lonza, Basel) with program FF-120. After the pulse, nucleofected cells were incubated for 10 minutes at 37°C and collected in a final volume of 200µl medium in a 96-well cell culture plate.

For the nuclefection of RNP into healthy donor (HD) or PKD-CD34^+^ cells, P3 Primary Cell 4D Nucleofector X Kit for Amaxa 4D device (Lonza, Basel, Switzerland) was used. Two hundred thousand cells were pre-stimulated for 24h or 48h and then resuspended in 20µl of nucleofection solution. RNP complex was added into the cellular suspension. The cells were nucleofected using DZ-100 program in strips. After the pulse, nucleofected HSPCs were incubated for 10 minutes at 37°C. Then, 180µl of pre-warmed medium was added and cells were transferred to a 96-well cell culture plate.

Nucleofected cells were immediately transduced with the corresponding AAV at different concentrations ranging from 10^4^ to 10^5^gc/µl in a 96-well culture plate in a final volume of 200 µL. Twenty-four hours after transduction, 100µl pre-warmed medium was added.

Gene editing in Jurkat cells was performed by transfection of RNP using the Nucleofector system (Lonza, Basel) as described previously *(49)*. In short, 5E5 cells were electroporated with RNPs using either Alt-R® S.p. Cas9 Nuclease V3 (IDT) or Alt-R® S.p. HiFi Cas9 Nuclease V3 at a concentration of 4 µM in the presence of 4.8 µM Alt-R® Cas9 Electroporation Enhancer (IDT). RNPs were generated by combining Cas9 and guideRNA complexes at a ratio of 1:1.2. GuideRNA complexes were generated by mixing Alt-R® CRISPR-Cas9 crRNA and Alt-R®CRISPR-Cas9 tracrRNA in equimolar amounts, followed by incubation for 5 minutes at 95°C and cooling to room temperature.

### Hematopoietic transplant protocol in immunodeficient mice

All the mice were kept under standard pathogen-free conditions in the animal facility of CIEMAT. All animal experiments were performed in compliance with European and Spanish legislations and institutional guidelines. The protocol was approved by “*Consejeria de Medio Ambiente y Ordenación del Territorio*” (Protocol number PROEX 073/15).

Human CD34^+^ cells were administered through tail vein of female NOD.Cg-*Prkdc*^*scid*^ *Il2rg*^*tm1Wjl*^/SzJ (NSG) or NOD.Cg-*Kit*^*W-41J*^*Tyr*^+^*Prkdc*^*scid*^*Il2rg*^*tm1Wjl*^/ThomJ (NBSGW) mice sub-lethally irradiated the day before transplant with 1.5Gy or 1Gy respectively. Together with these cells, 1×10^6^ irradiated with 20Gy hCD34^-^ cells (collected from the negative fraction of the CD34^+^ purification) were transplanted as support population. At day 30 after transplantation, bone marrow samples were obtained by intra-bone aspiration. NSG or NBSGW animals were euthanized at days 90 post-transplant or 60 post-transplant respectively. BM samples were collected and stained to analyze the percentage of gene-targeted cells.

To evaluate the long-term engraftment capacity of hCD34^+^ after AAV-mediated gene editing, the bone marrow cells extracted from primary NSG mice were transplanted into secondary NSG recipients. The human engraftment follow-up in secondary recipients was conducted as previously described, at 30 and 90 days post-transplantation.

### Flow Cytometry analysis

Flow cytometry analyses were conducted in LSR Fortessa Cell Analyser (BD/Becton Dickinson). Off-line analysis was made with FlowJo Software v10.5.0 (Tree Star).

To study the human HSPC phenotype of the gene edited cells were stained with hCD34-Pecy7 (581 clone, Biolegend), hCD38-PE (HB7 clone, BD) and hCD90-APC (5E10 clone, BD). After labelling, cells were washed and suspended in flow cytometry buffer containing 1µg/ml 4’,6-diamidino-2-phenylindole (DAPI, Thermo Fisher Scientific).

Human hematopoietic reconstitution in transplanted mice was analyzed by determining the percentage of progenitor cells (hCD34^+^), myeloid cells (hCD33^+^) and lymphoid cells (hCD19^+^) in the total engrafted human population subset (hCD45^+^). The BM cells were stained with hCD45-APCCy7 (HI30 clone, Biolegend), hCD34-APC (581 clone, BD), hCD19-PeCy7 (SL25C1 clone, Biolegend) and hCD33-PE (D3HL60.251 clone, Beckman Coulter). Afterwards, cells were washed and suspended in flow cytometry buffer containing 1µg/ml DAPI. Additionally, to assess human erythroid population within mouse BM, cells were stained with mTer119-FITC (TER119 clone, BioLegend), hCD235a-PE (HIR2 clone, BD) and hCD71-Pecy5 (M-A712 clone, BD) and suspended in flow cytometry buffer containing 1µg/ml DAPI.

### Genome targeting and quantification

To analyze the frequency of gene editing in HSPCs, specific integration of any of the two donors was assessed in colony forming units (CFUs). Two days after gene editing protocol, hCD34^+^ cells were resuspended in enriched methylcellulose medium (StemMACS™ HSC-CFU complete with Epo, Miltenyi Biotec). Fourteen days afterwards, colonies were counted and CFUs-GMs (granulocyte-macrophage colony forming units), BFU-Es (erythroid burst forming units) and CFU-GEMMs (granulocyte-erythroid-macrophage-megakaryocyte colony forming units) were identified and scored based on their morphological appearance in a blinded fashion. Individual colonies were picked in 100µl PBS. Genomic DNA from colonies was extracted by adding 20 µl of lysis buffer (0.3 mM Tris HCl pH 7.5, 0.6 mM CaCl2, 1,5 % Glycerol, 0.675 % Tween-20 and 0.3 mg/ml Proteinase K) and incubated at 65°C for 30 min, 90°C for 10 min and 4°C. After lysis, 30 µl of water were added, as previously described in Charrier et al *(50)*. Specific *in* and *out* primers [donorGFP-LHA/RHA and donorRPK-LHA/RHA] (see TableS5) were designed to amplify the regions of junction between the endogenous and exogenous DNA as shown in Figure S2 C. To assess the perfect insertion of the donor matrix, an in-out PCR was performed with Herculase II Fusion High-Fidelity DNA Polymerase (Agilent Technologies). The specific size of the PCR products was verified in 1% agarose gel. To analyze gene editing levels in cells engrafted in immunodeficient mice or erythroid differentiation experiments, cell DNA was purified using DNeasy Blood & Tissue Kit (QIAGEN) and the previously described in-out PCR strategy was performed.

### Expression analysis

The expression of coRPK mRNA was done by quantitative real time PCR (qRT-PCR). Firstly, mRNA from the edited cells was purified using TRIzol RNA Isolation kit (Thermo Fisher Scientific) according manufacturer’s instructions. The RNA was retrotranscribed to cDNA using the SuperScript VILO cDNA Synthesis Kit (Thermo Fisher Scientific). Finally, the cDNA was submitted to qRT-PCR using 7500 Fast Real-Time device and the Fast SYBR Green Master Mix (Applied Biosystems, Thermo Fisher Scientific). The primers for the amplification of the coRPK sequence were the following: coRPK F 5’-GGTGGTGCAGAAGATCGGAC-3’ and coRPK R 5’-GCAGATTCACGCCCTTTCTG-3’. GAPDH was used as housekeeping gene, and the quantification was done by comparison of the expression value of the transgene with the expression value of the GAPDH. The relative expression to GAPDH was calculated according Pfaffl’s method*(51)*. Additionally, the specific size of the PCR products was verified in 2% agarose gel.

### ATP quantification

To quantify ATP production, CellTiter-Glo® Luminescent Cell Viability Assay (Promega) was used. Human erythroid cells were plated in an opaque-walled 96-well plate. Control wells were prepared containing medium without cells to obtain a value for background luminescence. Then, the plate was equilibrated and incubated at room temperature for approximately 30 minutes. Afterwards, a volume of CellTiter-Glo® reagent equal to the volume of cell culture medium present in each well was added and cell solution was mixed for 2 minutes on an orbital shaker to induce cell lysis. In order to stabilize luminescent signal the plate was incubated at room temperature for 10 minutes. Finally, luminescence can be recorded using Genios Pro reader (Tecan).

## DATA AVAILABILITY

This study includes no data deposited in external repositories

## AUTHOR CONTRIBUTIONS

S.F.B. and O.Q.B. designed and performed the experiments and wrote the manuscript; O.A., R.S., M.D., I.O., helped in the experimental procedures; D.P.D. and J.C. contributed in the experimental design; M.S.S., R.T., M.A.B., performed and analyzed experiments; J.L.L.L. and P.B. provided samples, J.A.B. suggested procedures; M.P. contributed to the experiments design and suggested procedures; J.C.S designed the experiments, wrote the manuscript and provided grant support.

## CONFLIC OF INTEREST

J.C.S. and J.A.B. are consultants, hold shares and receive funding from Rocket Pharma. M.P. serves on the scientific advisory board of CRISPR Therapeutics and Graphite Bio. D.P.D. is employed by Graphite Bio. R.T., M.S.S. and M.A.B. are employed by Integrated DNA Technologies, Inc (IDT), which manufactures reagents similar to some described in the manuscript. R.T and M.A.B. own equity in DHR, the parent company of IDT. All other authors declare that no competing financial conflict exists.

## ACKNOWLEDGEMENTS

The authors would like to thank Miguel A. Martin for the careful maintenance of PKD deficient mice, and Mrs. Aurora de la Cal, María del Carmen Sánchez, Soledad Moreno, Nadia Abu-sabha, Montserrat Aldea and Mr. Sergio Losada for their dedicated administrative help. This work was supported by grants from “Ministerio de Economía, Comercio y Competitividad y Fondo Europeo de Desarrollo Regional (FEDER)” (SAF2017-84248-P), “Fondo de Investigaciones Sanitarias, Instituto de Salud Carlos III” (Red TERCEL; RD16/0011/0011) and Comunidad de Madrid (AvanCell, B2017/BMD-3692). The authors also thank Fundación Botín for promoting translational research at the Hematopoietic Innovative Therapies Division of the CIEMAT. CIBERER is an initiative of the “Instituto de Salud Carlos III” and “Fondo Europeo de Desarrollo Regional (FEDER)”.

## SUPPLEMENTARY MATERIALS

### Supplementary Figures

**Figure S1.**
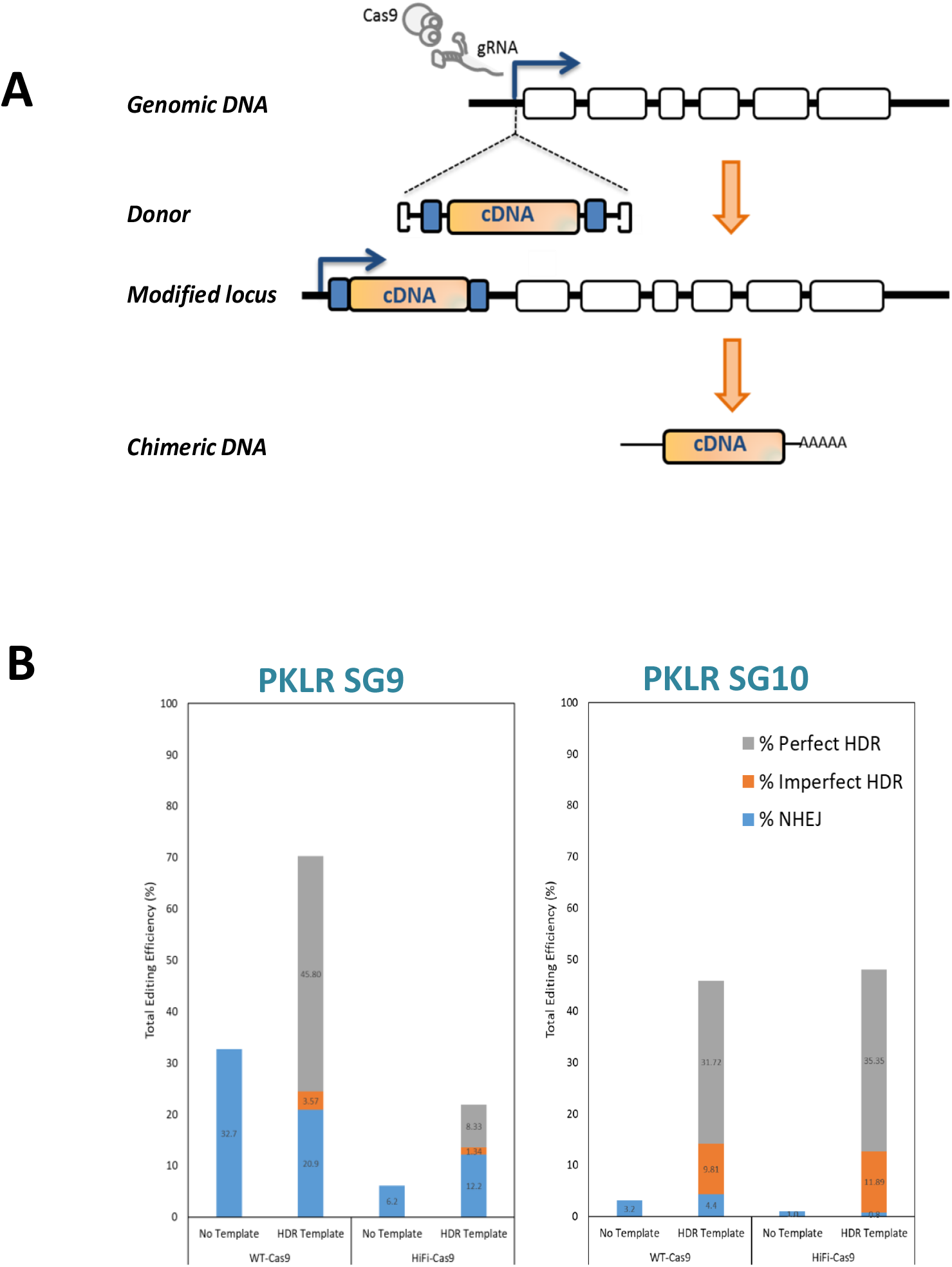
Design and optimization of the gene editing tools. **A)** Schematic of the gene editing protocol in the transcription starting site of *PKLR* gene. **B)** Representative TIDE analysis of SG1-RNP in CB-CD34+ cells. **C)** Identification of the off-targets produced by SG1-RNP in HSPCs by rhamp-seq. **D)** Analysis of the decomposition of the HDR in samples edited with SG4 and SG5 RNPs. Blue, % of NHEJ repair, Gray, % of perfect HDR, orange, % of imperfect HDR.

**Figure S2.**
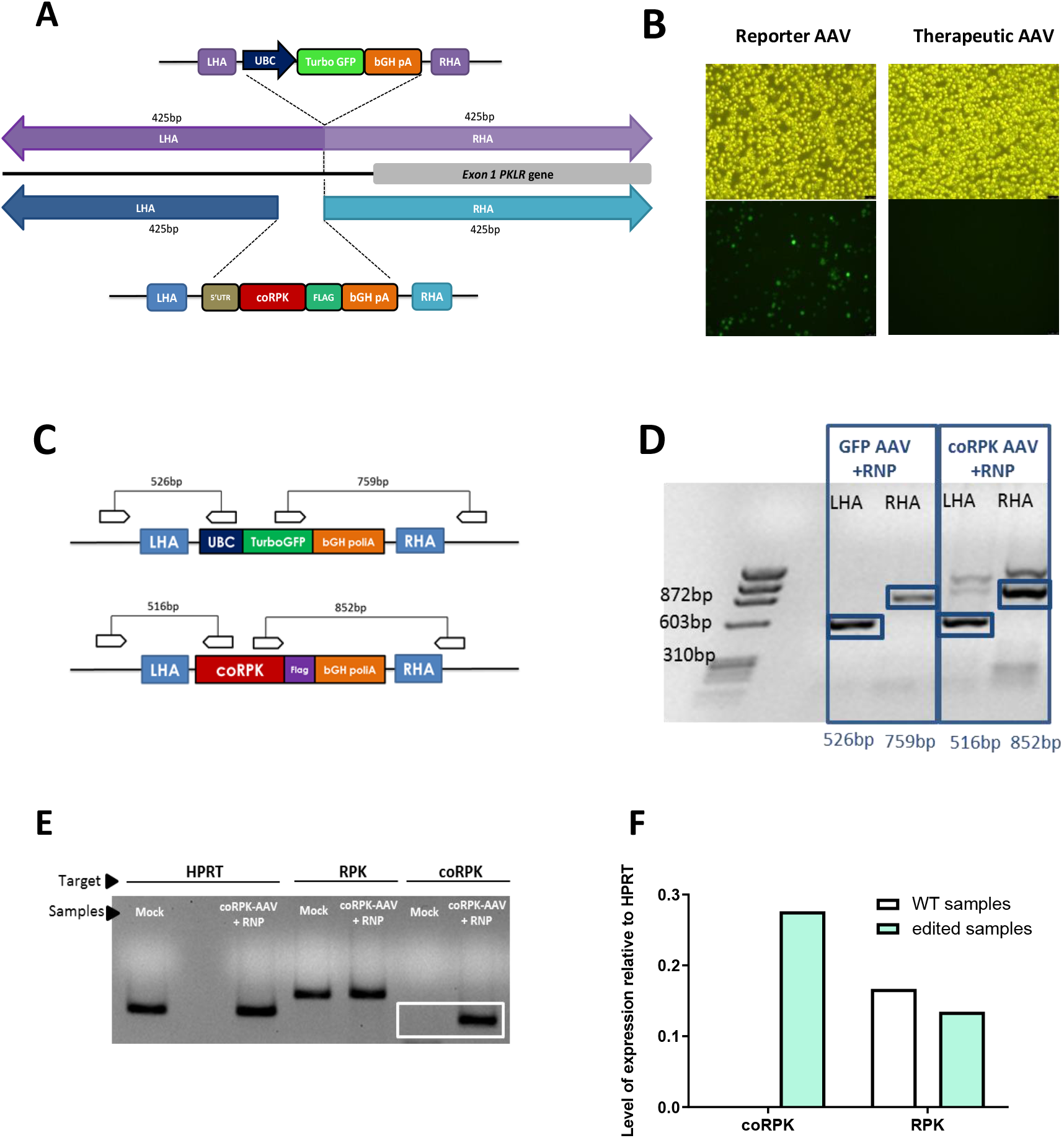
Gene editing in K562 cell line. **A)** Schematic representation of the HDR of reporter (up) and therapeutic donor (down). LHA: left homology arm; UBC: Ubiquitin C promoter; TurboGFP: Turbo green fluorescent protein; bGH pA: bovine growth hormone polyadenylation signal; RHA: right homology arm; 5’UTR: untranslated region; coRPK: codon optimized R-type Pyruvate Kinase; Flag: flag-tag sequence. **B**) Bright field and fluorecesce micrographs of K562 cells edited with either the AAV-GFP or the AAV-RPK donor vectors. Above 30% cells expressed GFP marker 5dpt. **C)** Specific in-out PCR strategy to assess the correct integration of both reporter and therapeutic donor at both 5’ (LHA) and 3’ (RHA) junctions. **D)** Specific bands verifying correct HDR in K562 cells. **E)** qRT-PCR products of samples transduced with therapeutic donor (coRPK-AAV+RNP) or untransduced. Housekeeping (HPRT), RPK and coRPK. F) Quantification of the qRT PCR. Levels of coRPK and RPK in mock (WT) and edited samples relativized to HPRT.

**Figure S3.**
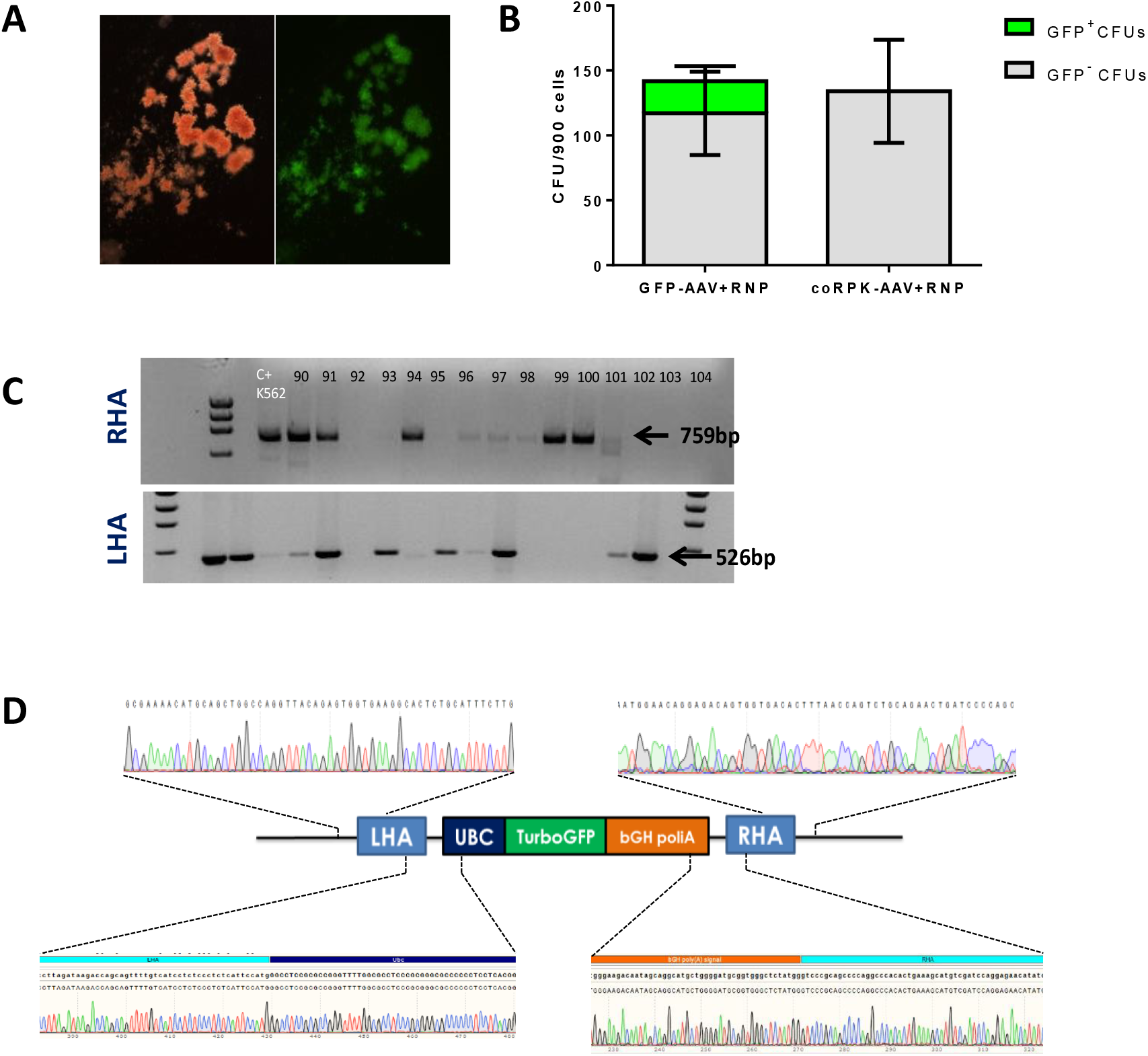
Gene editing using both donor AAVs in hematopoietic progenitors. **A)** Representative GFP^+^ BFU-E after gene editing using the AAV-GFP donor vector. **B)** Total number of CFUs and GFP^+^ CFUs after gene editing with either the AAV-GFP or the AAV-RPK donor vectors. Data are represented as mean ± SD; N=3 **C)** Representative agarose gel for the specific 5’ and 3’ PCRs in CFUs from cells edited with the AAV-GFP donor. **D)** Representative sanger sequencing of one of the positive amplicons

**Figure S4.**
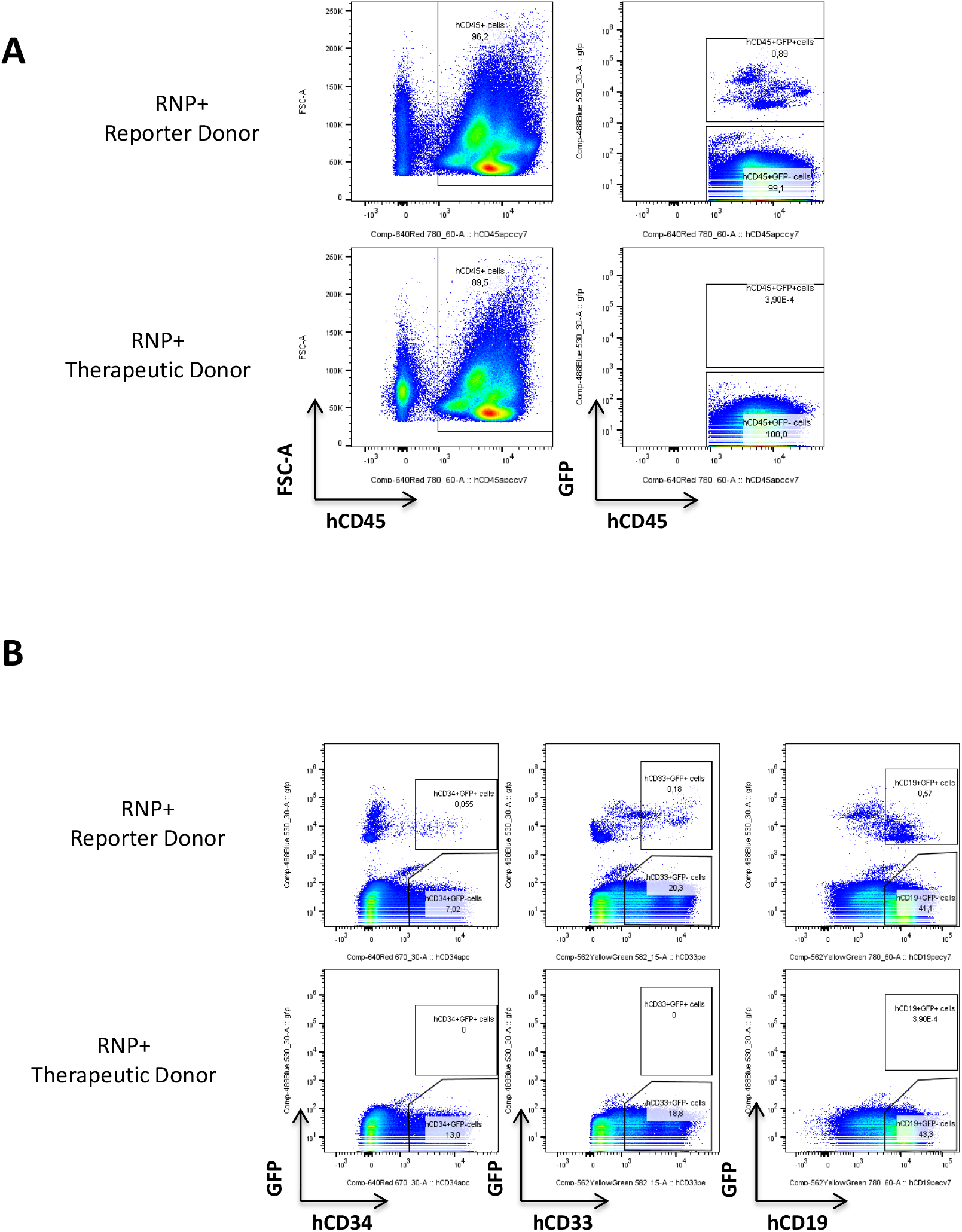
Representative flow cytometry dot-plots of human cell engraftment in immunodeficient NSG mice. **A)** FACS analyses of the BM of a mouse transplanted with HSPCs edited with the reporter donor (up) and from a mouse transplanted with cells edited with the therapeutic donor (bottom). **B)** Representative analyses of a multilineage engraftment by assessing CD34, CD33 and CD19 populations.

**Figure S5.**
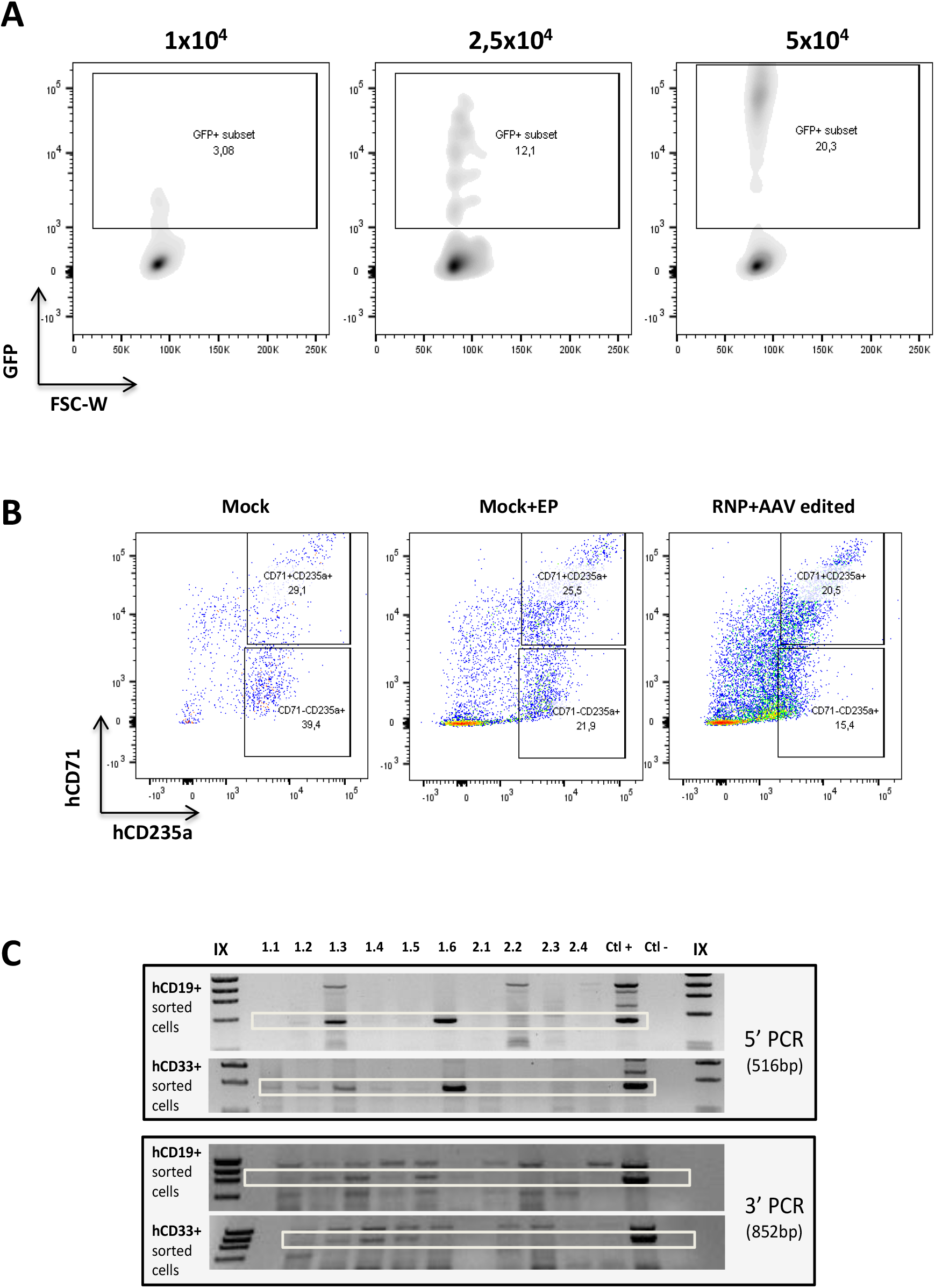
Optimizations in the gene editing protocol. **A)** FACS analysis of a representative sample of the reporter donor titration. **B)** Terminal erythroid differentiation of an edited PKD sample does not differ from the differentiation of unedited samples. **C)** 5’ and 3’ PCRs in hCD19+ and hCD33+ population sorted from the mice receiving edited cells (1.1, 1.2, 1.3, 1.4, 1.5 and 1.6) and from mice receiving unedited cells (2.1, 2.2, 2.3 and 2.4).

### Supplementary tables

**Table S1:**
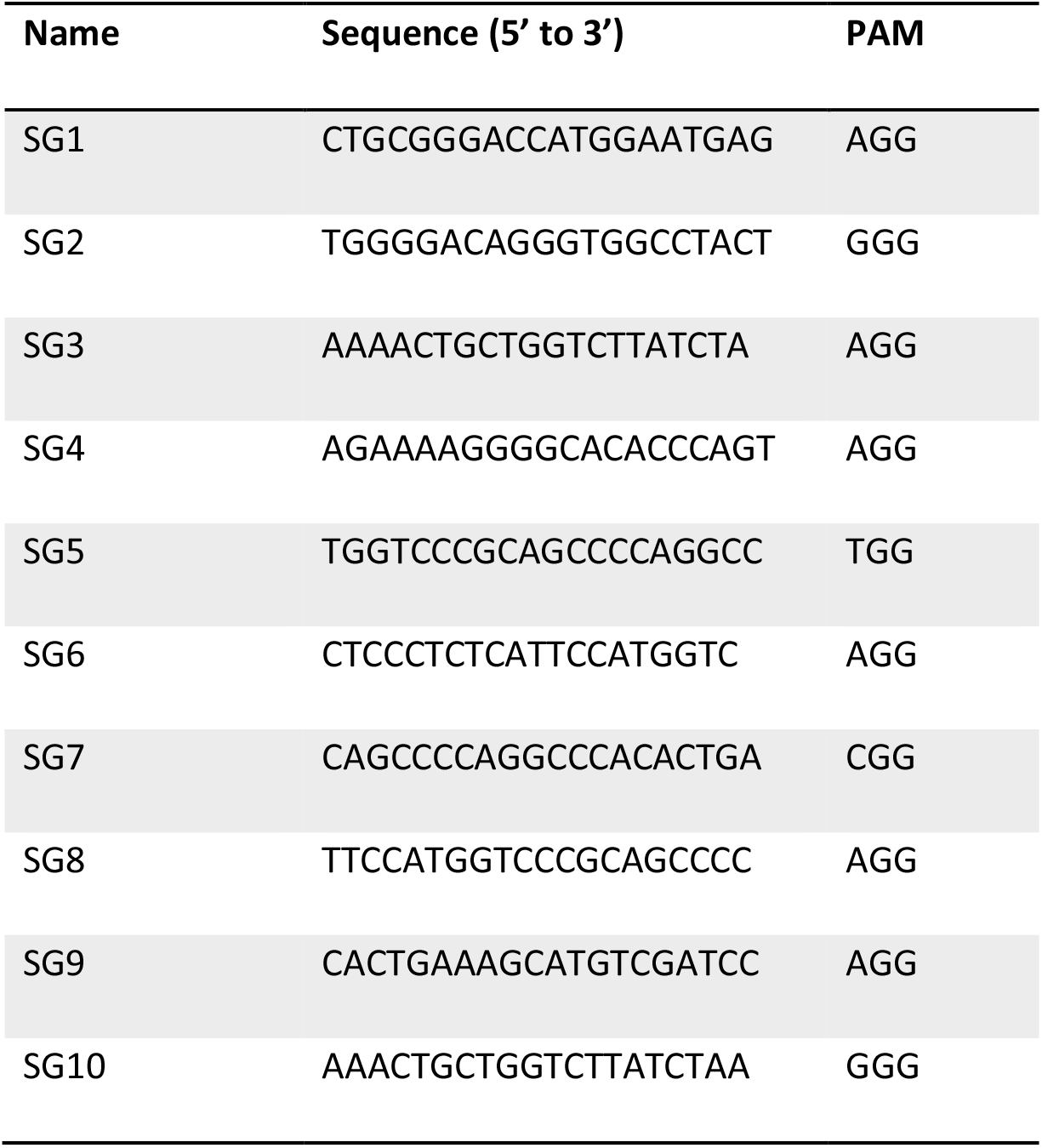
List of the different guide RNAs designed to target the transcription staring site of PKLR locus.

**Table S2:**
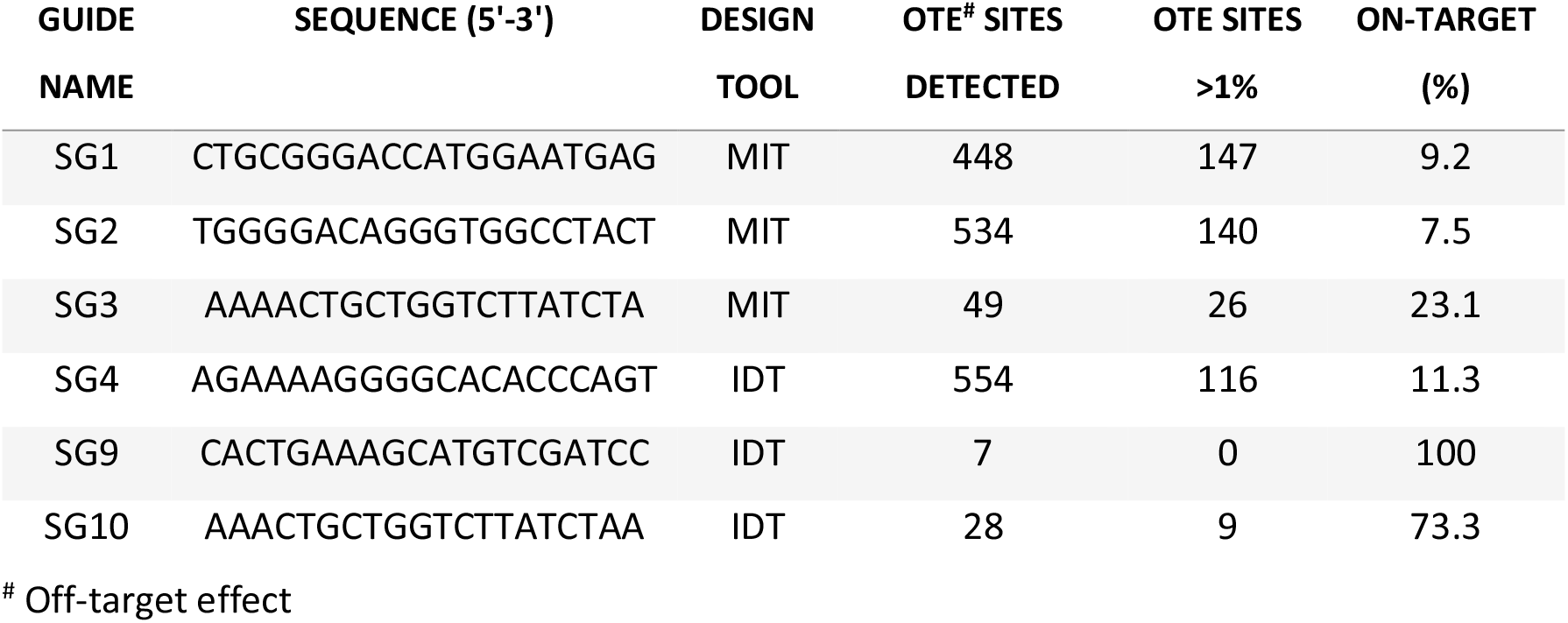
On-target and off-target analysis using GUIDE-Seq of the designed gRNAs in Hek293 cell line that constitutively expressed Cas9.

**Table S3:**
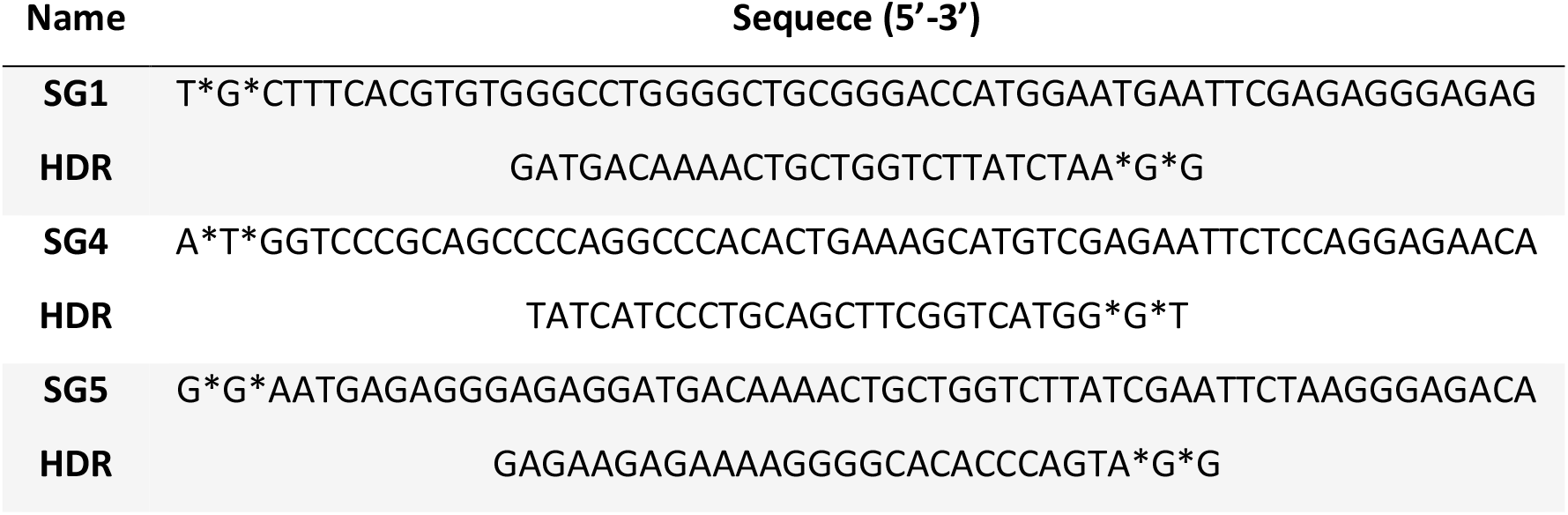
Sequence of the HDR templates used for the analysis of the gene editing nature.

**Table S4:** Specific mutations present in heterozygosity in the PKD samples used for the experiments.

| Patient | Mutations |
| --- | --- |
| **PKD-A** | c.1552C>A c.1456C>T |
| **PKD-B** | c.1003G>A c.1456C>T |
| **PKD-C** | c.721G>T c.1529G>A |
| **PKD-D** | c.359C>T c.1168G>A |

**Table S5:**
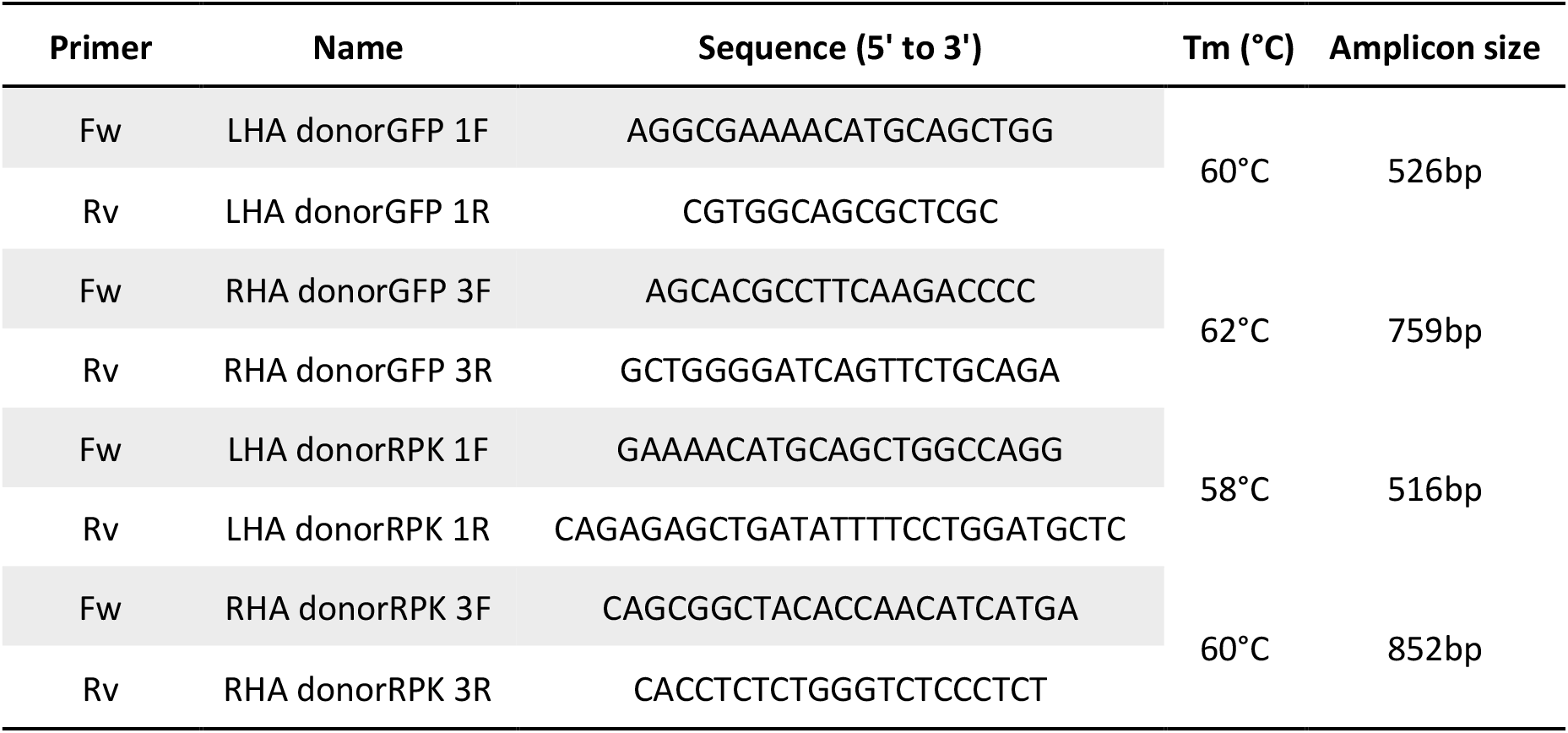
Primers used in the HDR analysis.

## Supplementary materials and methods

### *In vitro* erythroid differentiation

Human hematopoietic progenitors were differentiated towards erythroid lineage using a 14-day protocol, in which three different erythroid differentiation medium (EDM) were used. A basal medium was prepared and could be stored for 7 days at 4°C. Erythroid basal medium (EBM) was composed by IMDM, 2% FBS, 0.5% P/S solution, 3% human AB Serum (Sigma-Aldrich,), 10µg/ml Insulin (Sigma-Aldrich), 3U/ml Heparin (Sigma-Aldrich), and 200µg/ml Holo-transferrin (Sigma-Aldrich). Over basal medium, specific factors for the different steps of erythroid differentiation were added. Concentration was adjusted to 3×10^5^cells/ml for all the stages of the protocol. Cells were cultured in Erythroid Differentiation Medium-1 (EDM1) from days 0 to 6. EDM1 was prepared by adding 3U/ml EPO (Amgen), 10ng/ml SCF, 1ng/ml human interleukin 3 (hIL-3) (Eurobiosciences), and 1µM Dexamethasone (Sigma-Aldrich) to the EBM. On day 6, cell culture was transferred to Erythroid Differentiation Medium-2 (EDM2), composed by EBM supplemented with 3U/ml EPO and 10ng/ml SCF until day 9. For the last 5 days of the protocol, medium was based on EBM supplemented with 3U/ml EPO and increased concentration of holo-transferrin (200µg/ml) (EDM3). Cells were always cultured under normoxic conditions: 37°C, 21% O_2_, 5% CO_2_ and 95% relative humidity.

### rAAV cloning and production

A Reporter Donor was planned using SnapGene v1.1.3 software (GSL Biotech LLC). Homology Arms (HAs) of 425bp were designed around the SG1 guide RNA cutting site (see Table S1) in the *PKLR* starting site environment. The construct was comprised by 425bp-homologous arms (HAs) that flanked the Ubiquitin C Promoter (UBC) driving the expression of a Turbo-GFP sequence and a bGH polyadenilation signal (Figure S2). This donor sequence was cloned into a transfer plasmid carrying ITRs from pAAV-MCS plasmid (Agilent Technologies) containing AAV2 ITRs. The Therapeutic Donor, we aimed the expression of coRPK to be driven by the endogenous *PKLR* promoter, and thus, we did not add any promoter sequences. The RHA sequence was shared with Reporter Donor, but LHA was slightly different. Part of the 5’UTR sequence of *PKLR* gene was inserted together the codon optimized cDNA of RPK (coRPK, 1.7kb) *(19)*, a FLAG tag sequence, and the bGH polyadenylation signal (Figure S2).Consequently, the LHA was displaced 8bp upstream, leaving an 8-bp gap between the HAs corresponding to part of the 5’UTR sequence. The therapeutic donor was cloned similarly to reporter donor. Both therapeutic and reporter donors’ cDNA were diverged.

rAAV vectors were produced in-house or commercially by Vigene Biosciences. In brief, rAAVs were made in HEK 293T cells using standard PEI transfection with 6μ g ITR-containing plasmid and 22μg pDGM6 (kindly provided by Dr Russell), which contains the AAV6 cap genes, AAV2 rep genes, and adenovirus helper genes. After 72 hours, cells were harvested and centrifuged for 15 minutes 2000rpm. Cell pellet was resuspended in lysis buffer (2mM MgCl_2_ and 10mM TRIS/Cl, pH 8). Cells were lysed using five freeze-thaw cycles followed by 1h incubation at 37°C with Benzonase at 200 U/mL (Merck)). Finally, lysate was centrifuged at 4000rpm for 15 minutes at 4°C and the supernatant was collected. Purification of AAVs from the supernatant was conducted through iodixanol density gradient (Sigma-Aldrich) AAV vectors were extracted at the 58-40% iodixanol interface with 18-gauge metal syringe and stored at −80°C until the moment of use.

### Off-target analysis

Off-target analysis was performed based on the GUIDE-seq methodology *(52)*. The workflow was described previously *(40)*. In short, guideRNA complexes were generated using Alt-R® CRISPR-Cas9 crRNA XT and Alt-R®CRISPR-Cas9 tracrRNA, and transfected with a dsODN tag into HEK293 cells stably expressing Cas9 using the Nucleofector system (Lonza, Basel). After 72 hrs, genomic DNA was isolated followed by fragmentation and adapter ligation using the LOTUS™ DNA Library Prep Kit (IDT). Libraries were generated by two rounds of amplification, followed by Illumina-based NGS.

Targeted amplification of edited alleles was performed using the rhAmpSeq™ system (IDT), which consisted of two rounds of amplification. The first round employs locus specific primers, and the second round introduces Illumina-compatible P5/7 sequences to create dual-indexed amplicon libraries.

### Statistical analyses

Statistical analyses were conducted using Prism software (7.0 version, GraphPad). For the analyses of experiments in which n<5, a non-parametrical two-tailed Mann-Whitney test was performed when two variables were compared. For the comparison of multiple variables, Kruskal-Wallis with Dunn’s multiple comparison tests were performed. In experiments in which n>5, a Kolmogorov-Smirnov test was done to assess the normal distribution of the samples. If samples showed a normal distribution, a parametric two tailed paired t-test (when two variables were compared) or an ANOVA with Turkey’s multiple comparison tests when more variables were compared. If samples did not follow a normal distribution, the previously mentioned non-parametric tests were used. Significances and P values are indicated in the figures and/or legends. Additionally, the significance was represented by P-values: *P<0.05, **P<0.01, ***P<0.005 and ****P<0.001.

## REFERENCES

1. G. Canu, M. De Bonis, A. Minucci, E. Capoluongo, Red blood cell PK deficiency: An update of PK-LR gene mutation database, Blood Cells, Mol. Dis. 57, 100–109 (2016).

2. R. F. Grace, P. Bianchi, E. J. van Beers, S. W. Eber, B. Glader, H. M. Yaish, J. M. Despotovic, J. A. Rothman, M. Sharma, M. M. McNaull, E. Fermo, K. Lezon-Geyda, D. H. Morton, E. J. Neufeld, S. Chonat, N. Kollmar, C. M. Knoll, K. Kuo, J. L. Kwiatkowski, D. Pospíšilová, Y. D. Pastore, A. A. Thompson, P. E. Newburger, Y. Ravindranath, W. C. Wang, M. W. Wlodarski, H. Wang, S. Holzhauer, V. R. Breakey, J. Kunz, S. Sheth, M. J. Rose, H. A. Bradeen, N. Neu, D. Guo, H. Al- Sayegh, W. B. London, P. G. Gallagher, A. Zanella, W. Barcellini, Clinical spectrum of pyruvate kinase deficiency: Data from the pyruvate kinase deficiency natural history study, Blood 131, 2183–2192 (2018).

3. N. W. Meza, M. E. Alonso-Ferrero, S. Navarro, O. Quintana-Bustamante, A. Valeri, M. Garcia-Gomez, J. A. Bueren, J. M. Bautista, J. C. Segovia, Rescue of pyruvate kinase deficiency in mice by gene therapy using the human isoenzyme, Mol. Ther. 17, 2000–2009 (2009).

4. R. F. Grace, W. Barcellini, Management of pyruvate kinase deficiency in children and adults, Blood 136, 1241–1249 (2020).

5. R. Van Wijk, E. G. Huizinga, A. C. W. Van Wesel, B. A. Van Oirschot, M. A. Hadders, W. W. Van Solinge, Fifteen novel mutations in PKLR associated with pyruvate kinase (PK) deficiency: Structural implications of amino acid substitutions in PK, Hum. Mutat. 30, 446–453 (2009).

6. R. F. Grace, A. Zanella, E. J. Neufeld, D. H. Morton, S. Eber, H. Yaish, B. Glader, Erythrocyte pyruvate kinase deficiency: 2015 status report, Am. J. Hematol. 90, 825–830 (2015).

7. R. D. Christensen, L. D. Eggert, V. L. Baer, K. N. Smith, Pyruvate kinase deficiency as a cause of extreme hyperbilirubinemia in neonates from a polygamist community, J. Perinatol. 30, 233–236 (2010).

8. W. A. Muir, E. Beutler, C. Wasson, Erythrocyte pyruvate kinase deficiency in the Ohio Amish: Origin and characterization of the mutant enzyme, Am. J. Hum. Genet. 36, 634–639 (1984).

9. A. Zanella, E. Fermo, P. Bianchi, L. R. Chiarelli, G. Valentini, Pyruvate kinase deficiency: The genotype-phenotype association, Blood Rev. 21, 217–231 (2007).

10. I. Henig, T. Zuckerman, Hematopoietic Stem Cell Transplantation—50 Years of Evolution and Future Perspectives, Rambam Maimonides Med. J. 5, e0028 (2014).

11. S. E. Smetsers, F. J. Smiers, D. Bresters, M. C. Sonnevelt, M. B. Bierings, Four decades of stem cell transplantation for Fanconi anaemia in the Netherlands, Br. J. Haematol. 174, 952–961 (2016).

12. van W. R. van Straaten S, Bierings M, Bianchi P, Akiyoshi K, Kanno H, Serra IB, Chen J, Huang X, van Beers E, Ekwattanakit S, Güngör T Kors WA, Smiers F, Raymakers R, Yanez L, Sevilla J, van Solinge W, Segovia JC, Worldwide study of hematopoietic allogeneic stem cell transplantation in pyruvate kinase deficiency, (2018).

13. C. E. Dunbar, K. A. High, J. K. Joung, D. B. Kohn, K. Ozawa, M. Sadelain, REVIEW SUMMARY Gene therapy comes of age, Science 359, eaan4672 (2018).

14. J. A. Bueren, O. Quintana-Bustamante, E. Almarza, S. Navarro, P. Río, J. C. Segovia, G. Guenechea, Advances in the gene therapy of monogenic blood cell diseases, Clin. Genet. 97, 89–102 (2020).

15. A. A. Thompson, M. C. Walters, J. Kwiatkowski, J. E. J. Rasko, J. A. Ribeil, S. Hongeng, E. Magrin, G. J. Schiller, E. Payen, M. Semeraro, D. Moshous, F. Lefrere, H. Puy, P. Bourget, A. Magnani, L. Caccavelli, J. S. Diana, F. Suarez, F. Monpoux, V. Brousse, C. Poirot, C. Brouzes, J. F. Meritet, C. Pondarré, Y. Beuzard, S. Chrétien, T. Lefebvre, D. T. Teachey, U. Anurathapan, P. J. Ho, C. Von Kalle, M. Kletzel, E. Vichinsky, S. Soni, G. Veres, O. Negre, R. W. Ross, D. Davidson, A. Petrusich, L. Sandler, M. Asmal, O. Hermine, M. De Montalembert, S. Hacein-Bey-Abina, S. Blanche, P. Leboulch, M. Cavazzana, Gene therapy in patients with transfusion-dependent β-thalassemia, N. Engl. J. Med. 378, 1479–1493 (2018).

16. P. L. Marina Cavazzana-Calvo, Emmanuel Payen, Olivier Negre, Gary Wang, Kathleen Hehir, Floriane Fusil, Julian Down, Maria Denaro, Troy Brady, Karen Westerman, Resy Cavallesco, Beatrix Gillet-Legrand, Laure Caccavelli, Riccardo Sgarra, Leila Maouche-Chrétien, F, Transfusion independence and HMGA2 activation after gene therapy of human β-thalassaemia, 46, 564–574 (2011).

17. S. Marktel, S. Scaramuzza, M. P. Cicalese, F. Giglio, S. Galimberti, M. R. Lidonnici, V. Calbi, A. Assanelli, M. E. Bernardo, C. Rossi, A. Calabria, R. Milani, S. Gattillo, F. Benedicenti, G. Spinozzi, A. Aprile, A. Bergami, M. Casiraghi, G. Consiglieri, N. Masera, E. D’Angelo, N. Mirra, R. Origa, I. Tartaglione, S. Perrotta, R. Winter, M. Coppola, G. Viarengo, L. Santoleri, G. Graziadei, M. Gabaldo, M. G. Valsecchi, E. Montini, L. Naldini, M. D. Cappellini, F. Ciceri, A. Aiuti, G. Ferrari, Intrabone hematopoietic stem cell gene therapy for adult and pediatric patients affected by transfusion-dependent ß-thalassemia, Nat. Med. 25, 234–241 (2019).

18. J. A. Ribeil, S. Hacein-Bey-Abina, E. Payen, A. Magnani, M. Semeraro, E. Magrin, L. Caccavelli, B. Neven, P. Bourget, W. El Nemer, P. Bartolucci, L. Weber, H. Puy, J. F. Meritet, D. Grevent, Y. Beuzard, S. Chrétien, T. Lefebvre, R. W. Ross, O. Negre, G. Veres, L. Sandler, S. Soni, M. De Montalembert, S. Blanche, P. Leboulch, M. Cavazzana, Gene therapy in a patient with sickle cell disease, N. Engl. J. Med. 376, 848–855 (2017).

19. M. Garcia-Gomez, A. Calabria, M. Garcia-Bravo, F. Benedicenti, P. Kosinski, S. López-Manzaneda, C. Hill, M. Del Mar Mau-Pereira, M. A. Martín, I. Orman, J. L. Vives-Corrons, C. Kung, A. Schambach, S. Jin, J. A. Bueren, E. Montini, S. Navarro, J. C. Segovia, Safe and efficient gene therapy for pyruvate kinase deficiency, Mol. Ther. 24, 1187–1198 (2016).

20. J. L. López Lorenzo, S. Navarro, A. J. Shah, M. G. Roncarolo, J. Sevilla, L. Llanos, B. Pérez Camino de Gaisse, S. Sanchez, B. Glader, M. Chien, O. Quintana Bustamante, B. C. Beard, K. M. Law, M. Zeini, G. Choi, E. Nicoletti, J. A. Bueren, G. R. Rao, J. D. Schwartz, J. C. Segovia, Lentiviral Mediated Gene Therapy for Pyruvate Kinase Deficiency: A Global Phase 1 Study for Adult and Pediatric Patients, Blood 136, 47 (2020).

21. M. Jasin, Genetic manipulation of genomes with rare-cutting endonucleases, Trends Genet. 12, 224–228 (1996).

22. J. Bonafont, Á. Mencía, M. García, R. Torres, S. Rodríguez, M. Carretero, E. Chacón-Solano, S. Modamio-Høybjør, L. Marinas, C. León, M. J. Escamez, I. Hausser, M. Del Río, R. Murillas, F. Larcher, Clinically Relevant Correction of Recessive Dystrophic Epidermolysis Bullosa by Dual sgRNA CRISPR/Cas9-Mediated Gene Editing, Mol. Ther. 27, 986–998 (2019).

23. N. Zabaleta, M. Barberia, C. Martin-Higueras, N. Zapata-Linares, I. Betancor, S. Rodriguez, R. Martinez-Turrillas, L. Torella, A. Vales, C. Olagüe, A. Vilas-Zornoza, L. Castro-Labrador, D. Lara-Astiaso, F. Prosper, E. Salido, G. Gonzalez-Aseguinolaza, J. R. Rodriguez-Madoz, CRISPR/Cas9-mediated glycolate oxidase disruption is an efficacious and safe treatment for primary hyperoxaluria type I, Nat. Commun. 9, 1–9 (2018).

24. N. E. Bengtsson, J. K. Hall, G. L. Odom, M. P. Phelps, C. R. Andrus, R. D. Hawkins, S. D. Hauschka, J. R. Chamberlain, J. S. Chamberlain, Muscle-specific CRISPR/Cas9 dystrophin gene editing ameliorates pathophysiology in a mouse model for Duchenne muscular dystrophy, Nat. Commun. 8, 1–9 (2017).

25. P. Genovese, G. Schiroli, G. Escobar, T. Di Tomaso, C. Firrito, A. Calabria, D. Moi, R. Mazzieri, C. Bonini, M. C. Holmes, P. D. Gregory, M. Van Der Burg, B. Gentner, E. Montini, A. Lombardo, L. Naldini, Targeted genome editing in human repopulating haematopoietic stem cells, Nature 510, 235–240 (2014).

26. D. P. Dever, R. O. Bak, A. Reinisch, J. Camarena, G. Washington, C. E. Nicolas, M. Pavel-Dinu, N. Saxena, A. B. Wilkens, S. Mantri, N. Uchida, A. Hendel, A. Narla, R. Majeti, K. I. Weinberg, M. H. Porteus, CRISPR/Cas9 β-globin gene targeting in human haematopoietic stem cells, Nature 539, 384–389 (2016).

27. C. A. Vakulskas, D. P. Dever, G. R. Rettig, R. Turk, A. M. Jacobi, M. A. Collingwood, N. M. Bode, M. S. McNeill, S. Yan, J. Camarena, C. M. Lee, S. H. Park, V. Wiebking, R. O. Bak, N. Gomez-Ospina, M. Pavel-Dinu, W. Sun, G. Bao, M. H. Porteus, M. A. Behlke, A high-fidelity Cas9 mutant delivered as a ribonucleoprotein complex enables efficient gene editing in human hematopoietic stem and progenitor cells, Nat. Med. 24, 1216–1224 (2018).

28. S. S. De Ravin, L. Li, X. Wu, U. Choi, C. Allen, S. Koontz, J. Lee, N. Theobald-Whiting, J. Chu, M. Garofalo, C. Sweeney, L. Kardava, S. Moir, A. Viley, P. Natarajan, L. Su, D. Kuhns, K. A. Zarember, M. V. Peshwa, H. L. Malech, CRISPR-Cas9 gene repair of hematopoietic stem cells from patients with X-linked chronic granulomatous disease, Sci. Transl. Med. 9 (2017), doi:10.1126/scitranslmed.aah3480.

29. H. L. M. Suk See De Ravin, Andreas Reik, Pei-Qi Liu, Linhong Li, Xiaolin Wu, Ling Su, Castle Raley, Narda Theobald, Uimook Choi, Alexander H. Song, Andy Chan, Jocelynn R. Pearl, David E. Paschon, Janet Lee, Hannah Newcombe, Sherry Koontz, Colin Sweeney, David A. S, Targeted Gene Addition to a Safe Harbor locus in human CD34+ Hematopoietic Stem Cells for Correction of X-linked Chronic Granulomatous Disease, Nat. Biotechnol 88, 424–429 (2016).

30. B. Diez, P. Genovese, F. J. Roman-Rodriguez, L. Alvarez, G. Schiroli, L. Ugalde, S. Rodriguez-Perales, J. Sevilla, C. Diaz de Heredia, M. C. Holmes, A. Lombardo, L. Naldini, J. A. Bueren, P. Rio, Therapeutic gene editing in CD34 + hematopoietic progenitors from Fanconi anemia patients, EMBO Mol. Med. 9, 1574–1588 (2017).

31. P. Rio, R. Baños, A. Lombardo, O. Quintana-Bustamante, L. Alvarez, Z. Garate, P. Genovese, E. Almarza, A. Valeri, B. Díez, S. Navarro, Y. Torres, J. P. J. P. Trujillo, R. Murillas, J. C. J. C. Segovia, E. Samper, J. Surralles, P. D. P. D. Gregory, M. C. Holmes, L. Naldini, J. A. Bueren, O. Quintana-Bustamante, L. Alvarez, Z. Garate, P. Genovese, E. Almarza, A. Valeri, B. Díez, S. Navarro, Y. Torres, J. P. J. P. Trujillo, R. Murillas, J. C. J. C. Segovia, E. Samper, J. Surralles, P. D. P. D. Gregory, M. C. Holmes, L. Naldini, J. A. Bueren, Targeted gene therapy and cell reprogramming in Fanconi anemia, EMBO Mol. Med. 6, 835–848 (2014).

32. D. E. Bauer, S. C. Kamran, S. Lessard, J. Xu, Y. Fujiwara, C. Lin, Z. Shao, M. C. Canver, E. C. Smith, L. Pinello, J. Peter, J. Vierstra, R. A. Voit, G. Yuan, M. H. Porteus, A. John, G. Lettre, S. H. Orkin, A erythroid enhancer of BCL11A subject t genetic variation, 342, 253–257 (2014).

33. K. H. Chang, S. E. Smith, T. Sullivan, K. Chen, Q. Zhou, J. A. West, M. Liu, Y. Liu, B. F. Vieira, C. Sun, V. P. Hong, M. Zhang, X. Yang, A. Reik, F. D. Urnov, E. J. Rebar, M. C. Holmes, O. Danos, H. Jiang, S. Tan, Long-Term Engraftment and Fetal Globin Induction upon BCL11A Gene Editing in Bone-Marrow-Derived CD34+ Hematopoietic Stem and Progenitor Cells, Mol. Ther. - Methods Clin. Dev. 4, 137–148 (2017).

34. N. Psatha, A. Reik, S. Phelps, Y. Zhou, D. Dalas, E. Yannaki, D. N. Levasseur, F. D. Urnov, M. C. Holmes, T. Papayannopoulou, Disruption of the BCL11A Erythroid Enhancer Reactivates Fetal Hemoglobin in Erythroid Cells of Patients with β-Thalassemia Major, Mol. Ther. - Methods Clin. Dev. 10, 313–326 (2018).

35. Y. Wu, J. Zeng, B. P. Roscoe, P. Liu, Q. Yao, R. Lazzarotto, M. K. Clement, M. A. Cole, K. Luk, C. Baricordi, A. H. Shen, C. Ren, E. B. Esrick, J. P. Manis, D. M. Dorfman, A. Williams, A. Biffi, C. Brugnara, L. Biasco, C. Brendel, Highly efficient therapeutic gene editing of human hematopoietic stem cells, 25, 776–783 (2019).

36. H. Frangoul, D. Altshuler, M. D. Cappellini, Y.-S. Chen, J. Domm, B. K. Eustace, J. Foell, J. de la Fuente, S. Grupp, R. Handgretinger, T. W. Ho, A. Kattamis, A. Kernytsky, J. Lekstrom-Himes, A. M. Li, F. Locatelli, M. Y. Mapara, M. de Montalembert, D. Rondelli, A. Sharma, S. Sheth, S. Soni, M. H. Steinberg, D. Wall, A. Yen, S. Corbacioglu, CRISPR-Cas9 Gene Editing for Sickle Cell Disease and β-Thalassemia., N. Engl. J. Med. (2020), doi:10.1056/NEJMoa2031054.

37. Z. Garate, O. Quintana-Bustamante, A. M. Crane, E. Olivier, L. Poirot, R. Galetto, P. Kosinski, C. Hill, C. Kung, X. Agirre, I. Orman, L. Cerrato, O. Alberquilla, F. Rodriguez-Fornes, N. Fusaki, F. Garcia-Sanchez, T. M. Maia, M. L. Ribeiro, J. Sevilla, F. Prosper, S. Jin, J. Mountford, G. Guenechea, A. Gouble, J. A. Bueren, B. R. Davis, J. C. Segovia, Generation of a High Number of Healthy Erythroid Cells from Gene-Edited Pyruvate Kinase Deficiency Patient-Specific Induced Pluripotent Stem Cells, Stem Cell Reports 5, 1053–1066 (2015).

38. O. Q. Id, S. Fa, I. Orman, R. Torres, P. Duchateau, L. Poirot, J. A. Bueren, J. C. Segovia, Gene editing of PKLR gene in human hematopoietic progenitors through 5 ’ and 3 ’ UTR modified TALEN mRNA,, 1–20 (2019).

39. G. Schiroli, A. Conti, S. Ferrari, L. della Volpe, A. Jacob, L. Albano, S. Beretta, A. Calabria, V. Vavassori, P. Gasparini, E. Salataj, D. Ndiaye-Lobry, C. Brombin, J. Chaumeil, E. Montini, I. Merelli, P. Genovese, L. Naldini, R. Di Micco, Precise Gene Editing Preserves Hematopoietic Stem Cell Function following Transient p53-Mediated DNA Damage Response, Cell Stem Cell 24, 551-565.e8 (2019).

40. J. Shapiro, O. Iancu, A. M. Jacobi, M. S. McNeill, R. Turk, G. R. Rettig, I. Amit, A. Tovin-Recht, Z. Yakhini, M. A. Behlke, A. Hendel, Increasing CRISPR Efficiency and Measuring Its Specificity in HSPCs Using a Clinically Relevant System, Mol. Ther. - Methods Clin. Dev. 17, 1097–1107 (2020).

41. R. O. Bak, N. Gomez-Ospina, M. H. Porteus, Gene Editing on Center Stage, Trends Genet. 34, 600–611 (2018).

42. N. Hubbard, D. Hagin, K. Sommer, Y. Song, I. Khan, C. Clough, H. D. Ochs, D. J. Rawlings, A. M. Scharenberg, T. R. Torgerson, Targeted gene editing restores regulated CD40L function in X-linked hyper-IgM syndrome, Blood 127, 2513–2522 (2016).

43. R. A. Voit, A. Hendel, S. M. Pruett-Miller, M. H. Porteus, Nuclease-mediated gene editing by homologous recombination of the human globin locus, Nucleic Acids Res. 42, 1365–1378 (2014).

44. C. Lin, H. Li, M. Hao, D. Xiong, Y. Luo, C. Huang, Q. Yuan, J. Zhang, N. Xia, Increasing the Efficiency of CRISPR/Cas9-mediated Precise Genome Editing of HSV-1 Virus in Human Cells, Sci. Rep. 6, 1–13 (2016).

45. T. Maruyama, S. K. Dougan, M. Truttmann, A. M. Bilate, J. R. Ingram, H. L. Ploegh, Inhibition of non-homologous end joining increases the efficiency of CRISPR/Cas9-mediated precise [TM: inserted] genome editing, Nature 33, 538–542 (2015).

46. J. Song, D. Yang, J. Xu, T. Zhu, Y. E. Chen, J. Zhang, RS-1 enhances CRISPR/Cas9- and TALEN-mediated knock-in efficiency, Nat. Commun. 7, 1–7 (2016).

47. E. Haapaniemi, S. Botla, J. Persson, B. Schmierer, J. Taipale, CRISPR-Cas9 genome editing induces a p53-mediated DNA damage response, Nat. Med. 24, 927–930 (2018).

48. J. I. Garaycoechea, G. P. Crossan, F. Langevin, L. Mulderrig, S. Louzada, F. Yang, G. Guilbaud, N. Park, S. Roerink, S. Nik-Zainal, M. R. Stratton, K. J. Patel, Alcohol and endogenous aldehydes damage chromosomes and mutate stem cells, Nature 553, 171–177 (2018).

49. R. Basar, M. Daher, N. Uprety, E. Gokdemir, A. Alsuliman, E. Ensley, G. Ozcan, M. Mendt, M. H. Sanabria, L. N. Kerbauy, A. K. N. Cortes, L. Li, P. P. Banerjee, L. Muniz-Feliciano, S. Acharya, N. W. Fowlkes, J. Lu, S. Li, S. Mielke, M. Kaplan, V. Nandivada, M. Bdaiwi, A. D. Kontoyiannis, Y. Li, E. Liu, S. Ang, D. Marin, L. Brunetti, M. C. Gundry, R. Turk, M. S. Schubert, G. R. Rettig, M. S. McNeill, G. Kurgan, M. A. Behlke, R. Champlin, E. J. Shpall, K. Rezvani, Large-scale GMP-compliant CRISPR-Cas9-mediated deletion of the glucocorticoid receptor in multivirus-specific T cells, Blood Adv. 4, 3357–3367 (2020).

50. S. Charrier, M. Ferrand, M. Zerbato, G. Précigout, A. Viornery, S. Bucher-Laurent, S. Benkhelifa-Ziyyat, O. W. Merten, J. Perea, A. Galy, Quantification of lentiviral vector copy numbers in individual hematopoietic colony-forming cells shows vector dose-dependent effects on the frequency and level of transduction, Gene Ther. 18, 479–487 (2011).

51. Michael W. Pfaffl, A new mathematical model for relative quantification in real-time RT– PCR, Mon. Not. R. Astron. Soc. 29, 2003–2007 (2001).

52. S. Q. Tsai, Z. Zheng, N. T. Nguyen, M. Liebers, V. V Topkar, V. Thapar, N. Wyvekens, C. Khayter, A. J. Iafrate, L. P. Le, M. J. Aryee, J. K. Joung, GUIDE-seq enables genome-wide profiling of off-target cleavage by CRISPR-Cas nucleases, Nat. Biotechnol. 33, 187–197 (2015).

